# Structure of the mammalian ciliary transition zone microtubule doublet

**DOI:** 10.64898/2026.06.11.731632

**Authors:** Bin Cai, Aitor Pellicer Carmadiel, Emma J. van Grinsven, Timo van Veghel, Amol Aher, Ellen Aarts, Yixin Xu, Pedro Beltro, Jeffrey M. Beekman, Anna Akhmanova, Jingwei Xu, Michal Wieczorek

## Abstract

The ciliary transition zone gates bidirectional protein trafficking to maintain the specialized ciliary proteome using microtubule doublets as a scaffold. While ciliary axonemal doublets are well-characterized, the molecular architecture of the transition zone doublet remains elusive. Here, we report the structure of the mammalian transition zone doublet from bovine tracheal cilia using cryo-electron tomography at 4.7-5.0 Å resolution. The transition zone doublet is a structurally independent segment defined by an 8 nm-periodic arrangement of unique microtubule-inner proteins (MIPs) and microtubule-associated proteins (MAPs). We identify the calcium-binding protein CAPSL as a lumenal MIP that forms a pseudo-helical spiral and stabilizes microtubules *in vitro*. Furthermore, a dense MAP network on the A-microtubule surface clashes with intraflagellar transport (IFT) motor binding sites, suggesting anterograde IFT is directed to the B-microtubule for potential regulation by MAP9. Our work provides a structural framework for understanding gated ciliary transport and transition zone-linked human ciliopathies.

## Introduction

Cilia are hair-like organelles present in most eukaryotic lineages, with an evolutionary origin tracing back to the Last Eukaryotic Common Ancestor (LECA) (*1*). The assembly and maintenance of cilia depend on the bidirectional trafficking of protein complexes between the cilium and cytoplasm, which is carried out by intraflagellar transport (IFT) (*2*). To establish the highly specialized ciliary proteome (*3*), IFT trains must undergo stringent cargo checks before entering or exiting near the ciliary base via a region known as the transition zone, the molecular gatekeeper of the cilium (*4–7*).

The molecular basis of IFT cargo-checking by the transition zone is not clear. The transition zone is a ciliary segment located between the axoneme and the basal body (*8*). Some ultrastructural details of transition zones differ between species and cell types, including the presence of a so-called transition zone linker (TZ-linker) that connects neighboring doublets specifically in mammalian multiciliated cells (*8*). Nevertheless, a common feature in transition zones is the anchoring of microtubule doublets to the ciliary membrane by Y-shaped linkers, also called Y-links (*9*). One end of the Y-link binds to the outer junction between the A- and B-microtubule (AB outer junction) of the transition zone doublet, while the opposite end attaches to a membrane complex known as the ciliary necklace (*10*). In addition to the microtubule doublet, Y-link, and ciliary necklace, the transition zone contains transition fibers, which together form a loosely defined microtubule-membrane connection network. This network is thought to function as a selective filter that regulates the transport of both soluble and membrane proteins between the cytoplasm and the cilium (*7*), functionally analogous to the nuclear pore complex (*11*). Consequently, defective transition zones are linked to a variety of ciliopathies. For instance, mutations in previously identified proteins localized to the transition zone can cause Meckel-Gruber syndrome, nephronophthisis and Joubert syndrome (*12*).

Recent advances in cryo-EM single-particle analysis (SPA) have advanced our understanding of the ciliary axoneme and a few basal body components to high resolution in organisms including *Chlamydomonas*, human, and *Tetrahymena* (*13, 14*). Cryo-ET coupled with sub-tomogram averaging (STA) has similarly advanced our understanding of the ultrastructure of the basal body (*15*). Since IFT complexes are the active transporters that carry cargo into (anterograde) and out of (retrograde) the cilium, and the IFT motor protein complexes kinesin-2 (anterograde) and dynein-2 (retrograde) both bind to and walk along microtubules, IFT mobility depends on access to the surface of the transition zone doublet (*16, 17*). However, the molecular details of the transition zone, particularly its microtubule doublet, are poorly understood.

To elucidate the structural features of the transition zone, we employed a multi-scale imaging approach combining cryo-ET with STA, cryo-EM SPA, biochemical reconstitutions and fluorescence microscopy to generate a molecular model of the transition zone doublet from the bovine respiratory tract. We present cryo-ET reconstructions of the transition zone doublet at 4.7-5.0 Å resolution, which allow us to unambiguously assign its constituent microtubule inner proteins (MIPs) and microtubule associated proteins (MAPs). Our reconstructions reveal a doublet structure that is remarkably distinct from the axonemal doublet (*18*). We identified a previously uncharacterized MIP, calcyphosin-like (CAPSL), that heavily decorates the transition zone doublet lumen, consistent with an elevated expression level in tissues containing multiciliated cells. We also resolved a complex network of MAPs covering the outer surface of the doublet that almost entirely blocks the A-microtubule but leaves the B-microtubule accessible to IFT motors. We propose that this arrangement may direct anterograde IFT trains specifically to the B-microtubule, where they may be subjected to further regulation by MAP9, a protein implicated mainly in regulating intracellular trafficking in neurons (*19, 20*). Our structural model of the transition zone doublet from mammalian motile cilia reveals the structural basis of the transition zone scaffold and provides insights into its gatekeeping function via IFT regulation in motile cilia

## Results

### Cryo-ET structure of the bovine ciliary transition zone doublet

The transition zone is a weak point of the cilium (*21*); thus, mechanical isolation of cilia often causes breakage at the transition zone, leaving only the intact axoneme and basal body (*22*) (Fig. 1A). To obtain intact transition zones, we sought to isolate entire intact cilia using bovine trachea as our source material. The mammalian respiratory tract epithelium is composed of multiciliated cells, each with an apical membrane housing hundreds of embedded motile cilia (*23*), enabling us to isolate the apical membrane components (including entire cilia) through mild mechanical treatment in the presence of detergent (*22, 24*) (see Methods).

**Figure 1.**
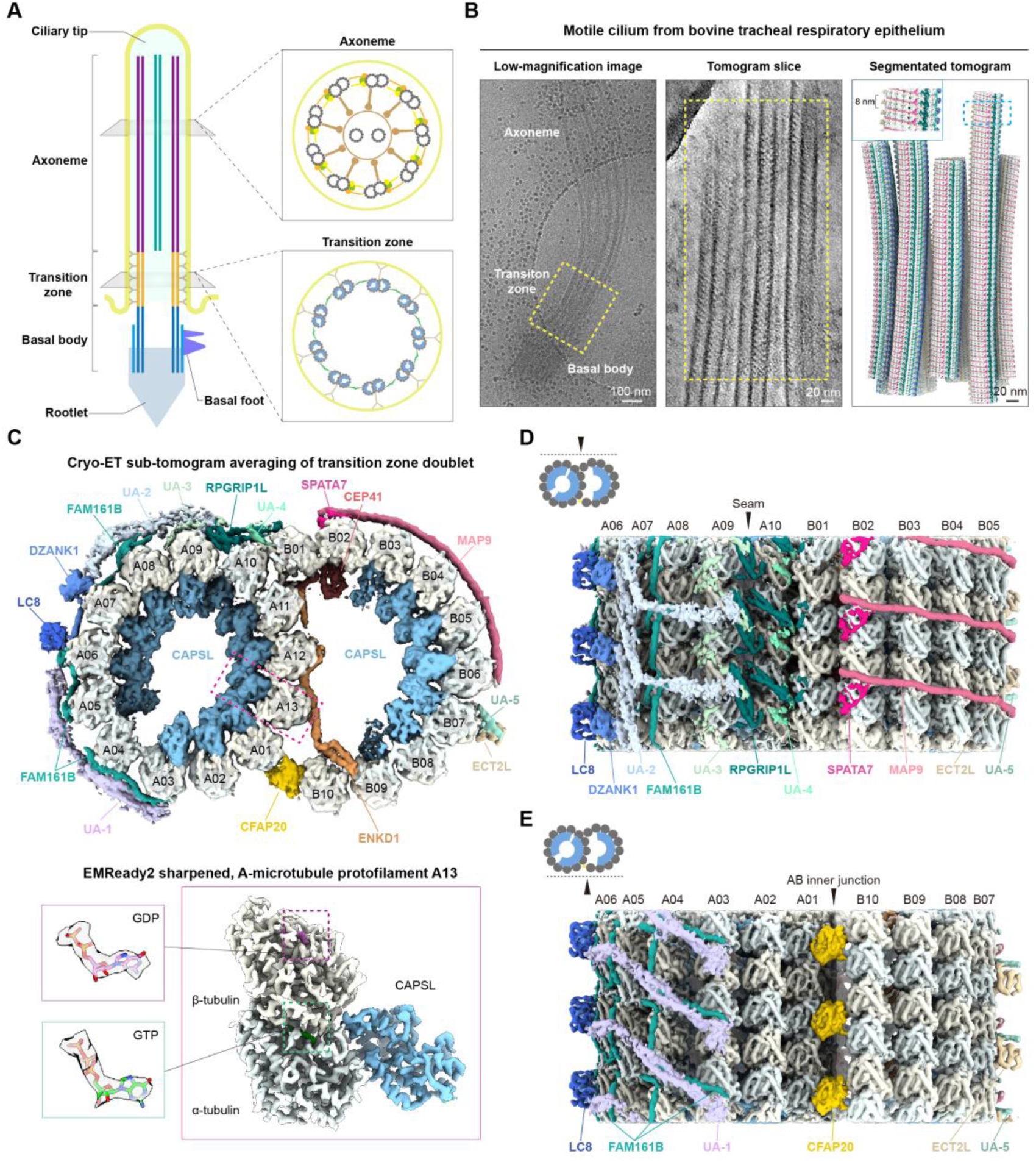
Cryo-ET structure of the bovine ciliary transition zone. **A.** Left: Schematic longitudinal cross section of a motile cilium. The ciliary tip, axoneme, transition zone, and basal body (including rootlet, distal appendage and basal foot) are indicated. Ciliary membrane (yellow), axonemal doublets (purple), central pair (cyan), transition zone doublets (orange), and basal body doublets/triplets (blue) are assigned different colors for clarity. Transition zone Y-links are shown to indicate attachments to the membrane and emphasize the transition zone as a gateway to the cilium. Insets on the right show cross section views of the axoneme and transition zone. Dynein complexes (axoneme) and Y-links, TZ-linkers, and MIPs (transition zone) are illustrated. **B.** Left: Low-magnification cryo-EM image of an intact cilium isolated from bovine trachea, with axoneme, transition zone (yellow dashed box), and basal body labeled. Midde: Slice through a cryo-tomogram (10 nm projection) of the transition zone, showing the region boxed on the left. Microtubule lumen striations are visible. Right: Segmented cryo-tomogram corresponding to the yellow dashed boxed region in the middle panel, generated using ArtiaX (*58*). Inset shows a zoom-in on the region in the blue dashed box highlighting the 8 nm periodicity of the transition zone doublet decorations. **C.** Upper: Cross-sectional view of the STA transition zone doublet composite map. Assigned protein densities are colored and labeled with protein names. Unassigned protein densities are colored and labeled with the prefix “UA”-1 through 5. Lower: Zoom-in view of the map within the pink dashed region in the upper panel, showing an EMReady2 (*59*) sharpened density of CAPSL bound to a tubulin dimer from A-microtubule protofilament A13. Insets show different nucleotide species (stick representations) in the tubulin heterodimer density (transparent surfaces). **D.** Outer surface view of the transition zone doublet composite map. Densities are colored and labeled as in panel **C**. The location of the A-microtubule seam is indicated. Unassigned protein densities are colored and labeled with the prefix “UA”-1 through 5. **E.** Inner surface view of the transition zone doublet composite map. Densities are colored and labeled as in panel **C**. The location of the AB inner junction is indicated. Unassigned protein densities are colored and labeled with the prefix “UA”-1 through 5.

Cryo-EM of vitrified cilia showed that all three major ciliary segments remained intact (Fig. 1B). The transition zone had a slightly narrower diameter compared to the basal body and the axoneme, which is a common feature of the transition zone in mammal cilia (*22, 25*). In our reconstructed tomograms, an 8-nm periodic pattern decorating the lumen of the transition zone doublets was clearly visible, and all transition zone microtubules appeared slightly bent inward, consistent with the narrower diameter of the entire transition zone (Fig. 1B).

Using cryo-ET and STA, we generated a 3D reconstruction of the transition zone microtubule doublet at 4.7-5.0 Å resolution from ∼1,300 tomograms (Fig.1C, fig. S1 and S2). The resulting local map quality was sufficient to distinguish GTP in α-tubulin vs. GDP in β-tubulin (Fig. 1C). The maps revealed an extensive network of MIPs and MAPs decorating the transition zone microtubule doublet with 8 nm periodicity (Fig. 1C, D and E). All but a few protein densities were identified using a combination of AI- and computational docking methods, including AlphaFold2/3 (*26, 27*), CryoAtom (*28*) and DomainSeeker (*29*). We assigned a total of 11 non-tubulin proteins to the transition zone doublet: calcyphosin-like (CAPSL), Enkurin domain-containing protein 1 (ENKD1), CEP41 and CFAP20 as MIPs; an RPGRIP1L dimer, FAM161B, a DZANK1 dimer, and a LC8 dimer as MAPs on the A-microtubule; and MAP9, SPATA7 and a dimer of ECT2L as MAPs on the B-microtubule (Fig. 1C, D and F>E, fig. S3, S4 and S5).

### MIPs in the transition zone doublet differ from those in the axonemal doublet

Previous studies of axonemal doublets have resolved a MIP network with over 29 proteins decorating the doublet lumen with 48 nm periodicity (*18*). In contrast, and despite being a physically connected segment of the axonemal doublet, the transition zone doublet exhibits a simpler MIP decoration with 8-nm periodicity (Fig. 2A). Of the 4 identified transition zone MIPs (CFAP20, CAPSL, ENKD1 and CEP41), only the AB inner junction protein CFAP20 is shared with the axoneme (Fig. 2B). The EF-hand domain-containing calcium-binding protein CAPSL is the predominant MIP in the transition zone doublet. CAPSL binds to α-tubulin of all protofilaments except A04, B01, B02 and B10, resulting in a series of spiral-shaped densities (Fig. 1C, Fig. 2A), consistent with previous lower resolution reconstructions (*17, 30*).

**Figure 2.**
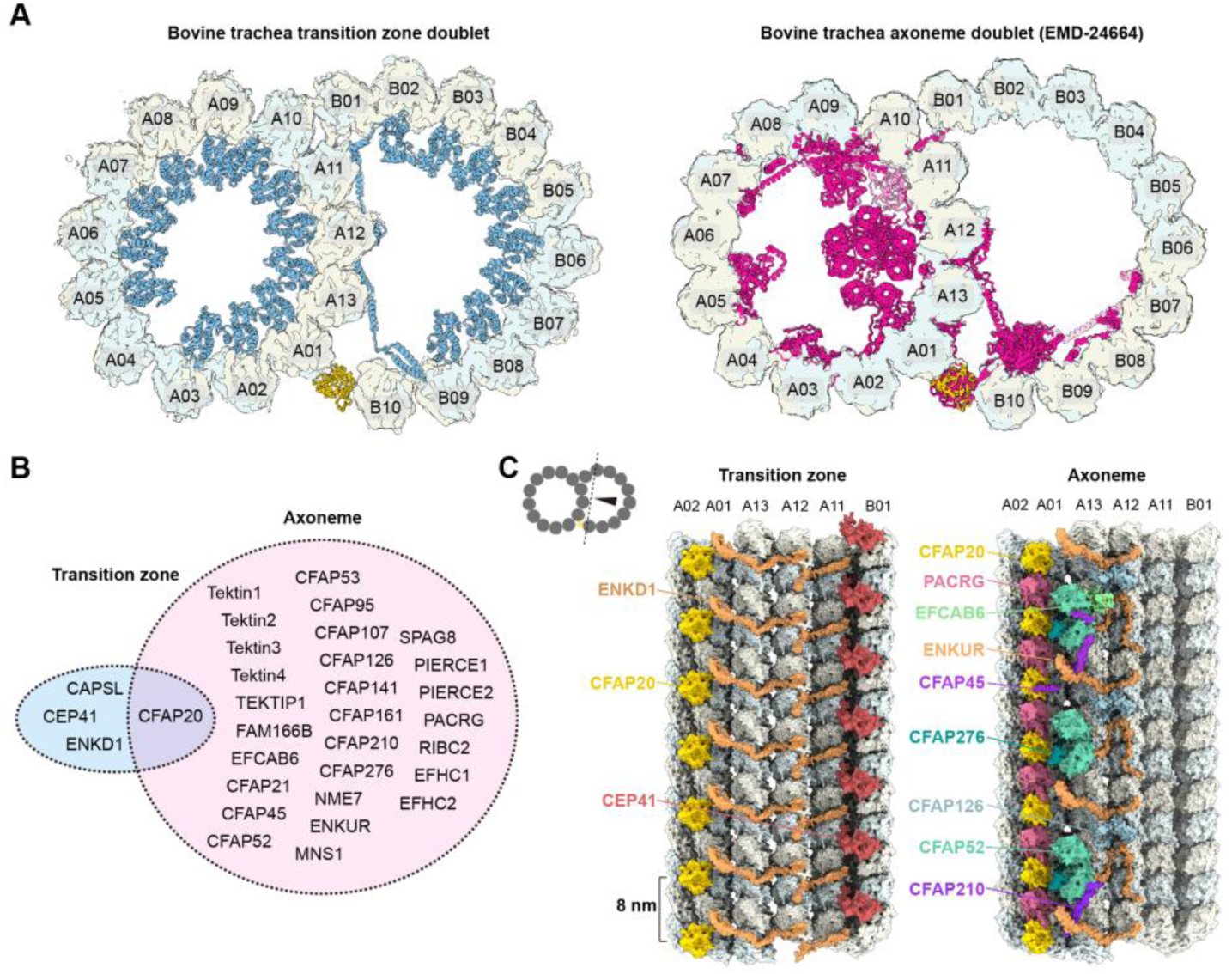
Comparison of MIPs in transition zone and axonemal doublets. **A.** Left: Cross section view of the transition zone doublet microtubule density map (transparent surface) superimposed with the MIP network (CAPSL, CEP41, and ENKD1; blue cartoon representation). CFAP20 is shown in yellow cartoon representation to contrast it with the other transition zone doublet MIPs. Right: Cross section view of the axonemal doublet microtubule density map (transparent surface) superimposed with the MIP network (pink cartoon representation). For comparison with the transition zone doublet, only CFAP20 is again shown in yellow cartoon representation to contrast it with the other axonemal doublet MIPs. **B.** List of MIPs shared between bovine trachea axonemal doublets (pink oval) (*18*) and the transition zone doublet (blue oval; this study). Only CFAP20 is shared between the two types of ciliary doublets. **C.** View of the transition zone doublet MIPs (left) and the axonemal doublet MIPs (right; (*18*)) on the A-microtubule surface as viewed from the B-microtubule lumen, showcasing the distinct MIP decoration pattern between the transition zone and axonemal doublets.

In addition to CFAP20 and CAPSL, both ENKD1 and CEP41 also decorate the B-microtubule lumen. ENKD1 is a paralog of Enkurin, a MIP found in the axonemal doublet (*18*). Whereas Enkurin and CFAP52 form a 16-nm repeating unit in the axoneme, ENKD1 spans from the A13 to the A11 protofilament to form an 8-nm repeating unit with CEP41 (Fig. 2C). These results show that the biochemical and spatial organization of most MIPs in the transition zone doublet of airway motile cilia are distinct from those in the axoneme.

### The transition zone MIP CAPSL is a microtubule-stabilizing protein

Several MIPs have been identified to form spiral-shaped structures analogous to CAPSL, including SPACA9, a MIP that decorates the lumen of mammalian sperm flagella tip microtubule singlets (*31–33*). Notably, CAPSL’s binding mode to α-tubulin clearly differs from SPACA9 and the previously reported MIPs that also form spiral densities in specialized microtubule lumens, such as Trx1 and Trx2 from *Toxoplasma gondii* (*34*), all of which instead engage with the intradimer and inter-protofilament interface of two neighboring α/β-tubulin subunits (fig. S5). To further validate the assignment of CAPSL to the spiral-shaped lumenal density in our STA reconstructions, as well as explore its potential effects on microtubule stability, we pursued three complementary approaches. First, we performed immunofluorescence staining of CAPSL in multiciliated cells differentiated from human nasal epithelial cells (*23, 35*). CAPSL antibody staining shows a clear dot-like pattern localized to the apical membrane near the base of cilia, as judged by tyrosinated tubulin staining, suggesting that CAPSL is a *bone fide* transition zone component (Fig. 3A, fig. S6A).

**Figure 3.**
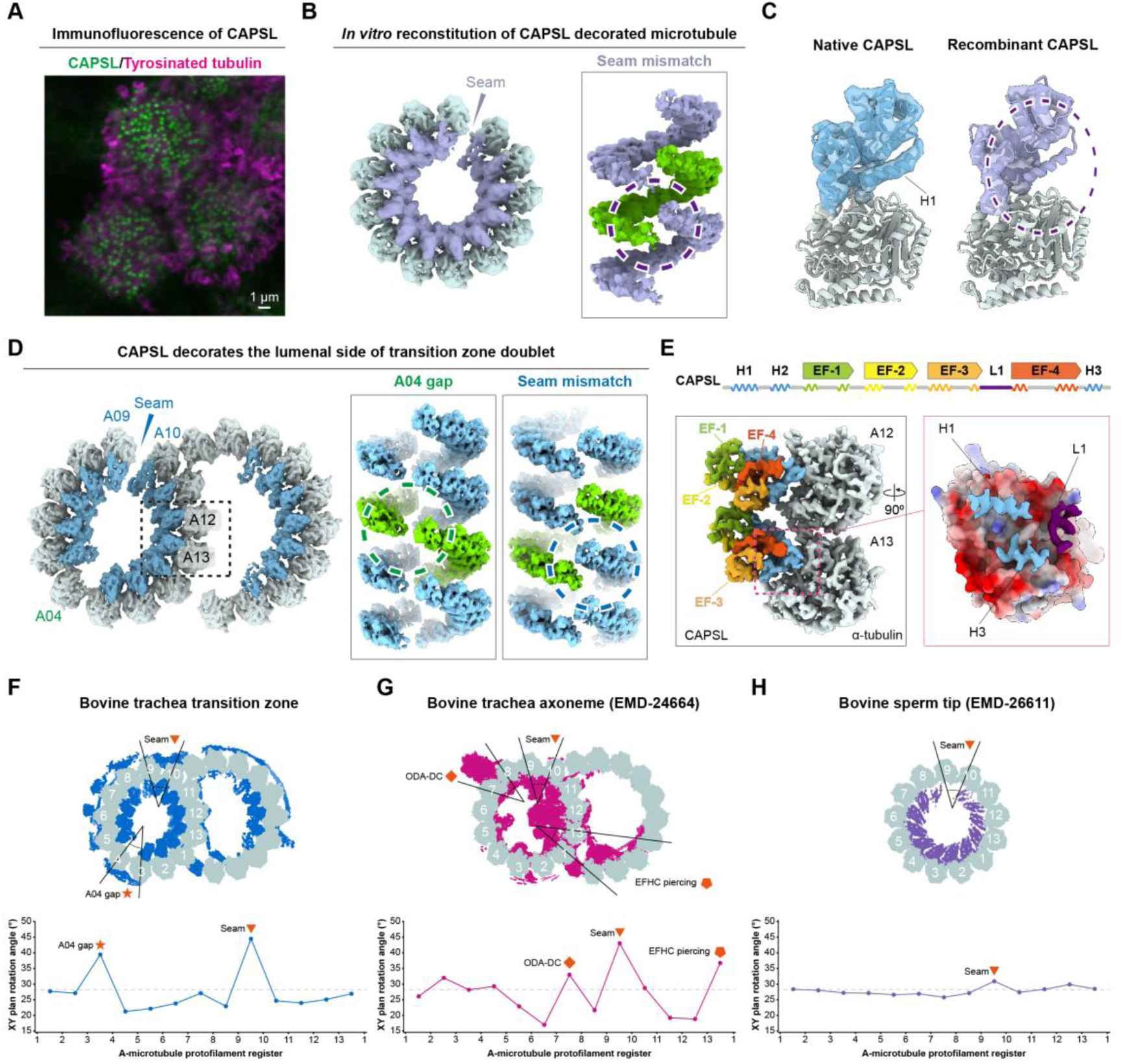
The microtubule-stabilizing MIP CAPSL forms a spiral-shaped structure in the lumen of transition zone doublet A- and B-microtubules. **A.** Merged immunofluorescence micrograph of the apical membrane of differentiated human airway multiciliated cells stained for CAPSL (green) and tyrosinated tubulin (magenta). **B.** Left: Cross section view of the single particle cryo-EM reconstruction of a paclitaxel-stabilized microtubule (pale blue) decorated with recombinant human CAPSL (light purple). Location of the seam is indicated. Right: Views of the segmented recombinant human CAPSL densities alone focusing on the seam and showing the lack of an A04 gap. Green surface indicates a single “turn” of the CAPSL spiral to highlight its continuity through the location in the reconstituted lattice that would correspond to the A04 gap. **C.** Comparison between the native bovine CAPSL model from the transition zone (left, light blue) and the corresponding model for recombinant human CAPSL (right, light purple) from the SPA reconstruction. CAPSL densities are shown in transparent surfaces for both structures to highlight that helix H1 density is not resolved in the recombinant human CAPSL reconstruction (dashed purple circle). **D.** Left: Cross section view of the transition zone doublet microtubule density map segmented to show tubulin (azure) and the CAPSL decoration (light blue). Locations for the seam and the A04 gap in the CAPSL decoration are indicated. Dashed box indicates region analyzed further in panel **E**. Right: Two opposing views of the segmented native bovine CAPSL densities alone focusing on the A04 gap or the seam. Green surface indicates a single “turn” of the CAPSL spiral to highlight its discontinuity in the A04 gap. **E.** Top: Schematic diagram depicting secondary structure (H1-H3 and L1) or folded domain (EF-1 to EF-4) elements of CAPSL. Bottom: Focused refinement map corresponding to the boxed region in panel **D** showing two neighboring α-tubulins from A12 and A13 (light grey) and the two associated CAPSL monomers, colored and labeled according to the schematic on top. Inset on the right shows a rotated view of one of the interfaces, with α-tubulin shown in surface representation colored according to electrostatic potential and CAPSL secondary structure binding elements shown as segmented densities colored according to the schematic on top. **F-H.** Comparison of per-protofilament X-Y plane rotation angles for the A-microtubules in bovine trachea transition zone doublets (**F**, this study; A04 gap and seam locations indicated), bovine trachea axonemal doublets (**G**, (*18*); ODA-DC, EFHC piercing and seam locations indicated) and microtubule singles in the tips of bovine sperm axonemes (**H**, (*31*); seam location indicated). For each structure, the microtubule lattice is schematized in light grey, and decorating MAPs or MIPs are colored as one solid color for clarity. Protofilament numbers are also indicated for reference.

Second, we purified full-length recombinant human CAPSL from *E. coli* and incubated it with paclitaxel-stabilized microtubules polymerized from porcine brain tubulin. SPA of the reconstituted CAPSL-microtubule complexes imaged using cryo-EM revealed that each α-tubulin in the 13-protofilament CAPSL-microtubule bound to CAPSL at a 1:1 stoichiometry, and the overall conformation of CAPSL matched that from the native bovine transition zone doublet (Fig. 3B, fig. S7). The main difference was that α-helix H1 of CAPSL was resolved in the native sample but not in the reconstituted sample, which may be due to the absence of post-translational modifications in tubulin or CAPSL in the reconstituted system, or architectural differences in the 13-protofilament microtubules in the two systems (described further below) (Fig. 3C).

Third, and to test the effect of CAPSL-binding on microtubule stability, we visualized the effect of CAPSL on microtubule assembly *in vitro* using total internal reflection fluorescence (TIRF) microscopy. The addition of CAPSL significantly increased the number of microtubules nucleated by human γ-tubulin ring complexes, suggesting CAPSL stabilizes microtubules (fig. S8) (*36*).

### MAP decoration likely induces CAPSL spiral breakage

Interestingly, CAPSL formed a continuous spiral in the *in vitro* reconstituted microtubule, with only one mismatch at the seam where α-tubulin laterally interacts with β-tubulin (Fig. 3B). In contrast, the CAPSL spiral in the native transition zone doublet displays an additional break at the A04 protofilament (Fig. 3D). To understand the cause of the break at A04, we analyzed the CAPSL-α-tubulin interface in the native transition zone doublet. The four calcium-binding EF-hands of CAPSL point away from tubulin, while the H1, H3 helices and L1 loop are bound to the lumenal surface of α-tubulin. Two neighboring CAPSL molecules interact via the EF-1 domain of one CAPSL and the EF-3 domain of the adjacent CAPSL (Fig. 3E). At the A04 protofilament, the CAPSL binding site is exposed, but CAPSL is persistently vacant. This suggests that the break likely arises from geometrical restrictions in the microtubule lattice that preclude lateral CAPSL:CAPSL interactions from forming here.

To quantify the geometrical distortion of the microtubule lattice, we measured the relative angle between neighboring α-tubulin molecules in the A-microtubule, with deviations from the average indicating local distortions in protofilament-protofilament lateral contacts. Comparative analysis of the A-microtubule from our transition zone doublet (Fig. 3F), the *in vitro* reconstituted CAPSL-microtubule (fig. S9), the axonemal doublet (*18*) (Fig. 3G), and the SPACA9-decorated sperm tip singlet (*31*) (Fig. 3H) revealed marked distortion near the A04 protofilament only in the transition zone doublet reconstruction. As the A04 protofilament has no MIPs bound to it, the distortion is likely introduced by transition zone MAPs. Notably, FAM161B binds along the A04 protofilament and forms a tri-helical bundle with an unidentified coiled-coil density that stretches across the A03–A04 protofilament (fig. S4B).

The presence of this bundle likely widens the A03–A04 distance, stabilizing the distortion of the A-microtubule and consequently breaking the CAPSL spiral-shaped decorations in the lumen.

### Transition zone MAPs present a steric block to IFT motor proteins in the A-microtubule

In our STA reconstructions of the transition zone doublet, we observed a dense network of MAPs covering the outer surface of A-microtubule protofilaments A07-A10 (Fig. 4A). Additionally, the Y-link has been reported to dock near the A09-A10 (the seam of A-microtubule) protofilaments of the transition zone doublet (*10*); consistent with this, we identified RPGRIP1L, a previously-proposed Y-link protein component (*7, 37*), as a microtubule binding MAP whose N-terminal α-helices dimerize into coiled-coil segments that span precisely across protofilaments A09-A10 (fig. S4C). Together, the MAPs and the Y-link (through RPGRIP1L) form a rigid meshwork that occupies nearly the entire surface of the A-microtubule. Notably, this meshwork presents a steric clash with predicted microtubule-bound kinesin-2 or dynein-2 complexes (which drive anterograde and retrograde IFT, respectively) docked into most available motor binding sites on the A-microtubule (fig. S10). This suggests that motor proteins, including those responsible for IFT, would not be able to easily translocate across the transition zone via the A-microtubule.

**Figure 4.**
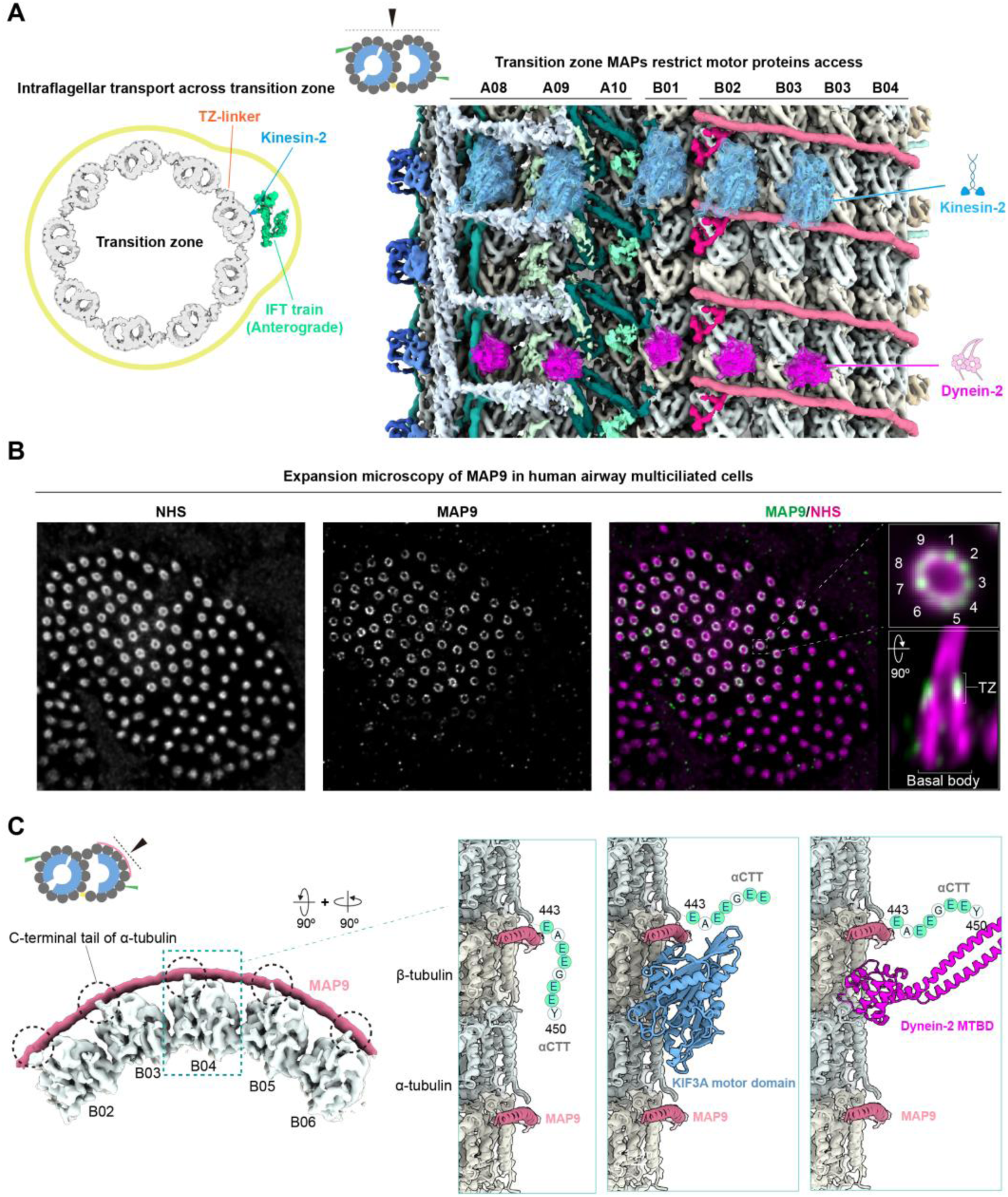
Transition zone MAPs block access of IFT motor proteins to the A-microtubule and may regulate IFT motor proteins on the B-microtubule. **A.** Left: Schematic depicting the relative size of an anterograde IFT train complex (green) carried by a motor protein (blue) relative to the microtubule doublet network in the transition zone. The ciliary membrane (yellow) is indicated and shown as a bulge to highlight the tight spacing for IFT transport across the transition zone. The location of the TZ-linker relative to the IFT train is also indicated. Right: View of the transition zone STA map from the outer surface (see schematic in top left), showing the dense network of MAPs obstructing the microtubule lattice in the A-microtubule (subunit densities colored as in Figure 1). Predicted complexes between tubulin and the kinesin-2 (KIF3A) motor domain (blue) or dynein-2’s microtubule binding domain (magenta) are shown docked into various A- and B-microtubule protofilaments and are displayed as cartoon representations overlaid with transparent surface representations. **B.** Ten-Fold Robust Expansion (TREx) microscopy micrograph of the cortex of a differentiated human airway multiciliated cell stained for whole-protein (NHS, left) and MAP9 (middle). A merge of the two channels is shown on the right, with insets showing cross sectional (top) and transverse longitudinal (bottom) views of a single cilium. The 9-fold symmetry of the MAP9 staining and its localization to the transition zone relative to the basal body are indicated. **C.** Left: Cross section view (see inset on top left) of the transition zone doublet map segmented to show B-microtubule protofilaments B02-B06 and the associated MAP9 density. Black dashed circles indicated protruding density for the C-terminal tails of adjacent α-tubulins contacting MAP9. Right: Series of the same views of two repeats of MAP9 (pink cartoon representation) bound to β-tubulin in the transition zone reconstruction, showcasing how the microtubule-binding domains of the ciliary motor proteins KIF3A (blue cartoon representation from an AlphaFold prediction) or dynein-2 (magenta cartoon representation) should generally be permitted to associate with the MAP9-decorated lattice. Schematics for the residues corresponding to the α-tubulin C-terminal tails associated with MAP9 and potentially position to further interact with different motors in a PTM-dependent manner (glutamates, colored in green) are indicated.

### MAP9 decorates a large portion of the outer surface of the B-microtubule

In contrast to the highly decorated A-microtubule, the B-microtubule is more sparsely decorated by: MAP9, SPATA7 and an ECT2L dimer. Only a small segment of SPATA7 binds to protofilament B02, and a similarly small segment of the ECT2L dimer binds to protofilament B07 as one end of the TZ-linker (Fig. 1D and E). In contrast, MAP9 spans across B-microtubule protofilaments B02-B06, and thus covers most of the B-microtubule’s motor-accessible surface (Fig. 1D). MAP9 is a highly conserved protein that stabilizes microtubules and selectively regulates kinesin and dynein motors. It has previously been shown to function in neurons, mitotic spindles and primary cilia (*38–43*), but whether it also functions in the motile cilia of multiciliated cells has not been fully explored.

To further validate our assignment of MAP9 to the transition zone, we performed expansion microscopy using a MAP9 antibody in multiciliated cells differentiated from human nasal epithelial cells (*35*); cilia were identified by fluorescent-NHS staining. We observed MAP9 signal mapped specifically to the transition zone region. The cross-sectional view revealed a nine-fold symmetrical distribution of the MAP9 signal (Fig. 4B), confirming MAP9 as a conserved transition zone component in mammalian motile cilia (*41*).

Docking of kinesin-2 and dynein-2 complexes into our transition zone B-microtubule revealed no obvious clash between MAP9 and the kinesin-2 and dynein-2 microtubule-binding domains (Fig. 4A, fig. S10). However, part of loop L8 of the kinesin-2 motor domain is predicted to come into close contact with the MAP9 helix (fig. S10). This loop has been implicated in regulating kinesin motor activity on MAP9-decorated microtubules *in vitro* (*43*), suggesting MAP9 may be involved in differential regulation of motor protein activity in motile cilia.

Although neither kinesin-2 nor dynein-2 motor proteins were resolved in our reconstructions, we note that density for the C-terminal tail of α-tubulin also extends towards MAP9 (Fig. 4C). This is consistent with a recent cryo-EM SPA structure of an *in vitro* reconstituted MAP9-microtubule complex, which suggested that MAP9 motifs that bind to β-tubulin are positively charged and attract the negatively charged α-tubulin tail (*43*), bringing the tubulin tail into proximity with motor proteins to potentially regulate motor activity via ciliary tubulin post-translational modifications (*19, 20*). Notably, the B01 protofilament at the AB outer junction does not present clashes with either kinesin-2 or dynein-2 (fig. S10), suggesting this lone protofilament could also serve as a high-affinity track for both anterograde and retrograde IFT motors.

## Discussion

Our results demonstrate that the transition zone microtubules are structurally distinct from those in both the axoneme and the basal body. While the axonemal doublet exhibits well-defined 48-nm internal and 96-nm external periodicities defined by specific MIPs and MAPs (*13, 18*), these patterns are absent in the 8-nm periodicity of the transition zone doublet. Furthermore, although the basal body triplet reportedly displays 8-nm or 16-nm periodicity in lower-resolution cryo-ET reconstructions (*15, 44–47*), its MIP densities do not share obvious similarity with those observed in the transition zone. Together, these findings indicate that the architecture of the transition zone is not dictated by the axoneme or basal body but rather constitutes a biochemically independent segment of the cilium.

We identified the calcium-binding protein CAPSL as the primary MIP within the transition zone microtubule doublet. To determine whether CAPSL is a conserved component across diverse ciliary types, we analyzed its abundance at the protein (*48*) and transcriptional levels (*49*). These data indicate markedly elevated CAPSL expression in human brain, respiratory organs, and reproductive systems (fig. S11A). Immunohistochemistry further confirms CAPSL enrichment at the apical membrane of multiciliated cells in human respiratory, efferent duct, and fallopian tube epithelia (fig. S11B). These data suggest that CAPSL is a ubiquitous component of transition zones in multiciliated cells from different tissues.

The epithelial cells in these tissues possess motile cilia that beat synchronously to propel mucus or gametes (*50*). This constant beating induces mechanical stress that could challenge ciliary structural integrity. In motile cilia, the axoneme is reinforced by dynein complexes and a central apparatus (*13*), while the basal body is supported by the rootlet (*51*); these features are notably absent or reduced in non-motile cilia (*13*). Thus, compared to the basal body, which is additionally supported by the apical membrane-cytoskeleton network (*17, 52*), the transition zone likely constitutes a mechanical weak point that must withstand the shear forces from ciliary beating (*21*). Given its elevated expression in tissues containing multiciliated cells and our *in vitro* analysis on recombinant CAPSL, we propose that part of CAPSL’s function is to stabilize the transition zone doublet. Beyond CAPSL, the MAPs densely coating the transition zone doublet may further contribute to structural stability. For instance, the microtubule-binding helix of MAP9 resolved in our structure is known to stabilize microtubules against depolymerization (*43*).

The specialized architecture of the transition zone membrane-microtubule network comprising the transition zone doublet, Y-links, and ciliary necklace, suggests a possible mechanism for the asymmetric regulation of IFT. Our structure reveals that several MAPs including the Y-link docking protein RPGRIP1L should broadly restrict motor protein access to the A-microtubule. Conversely, the B02-B06 protofilaments are decorated by MAP9 and SPATA7 in a manner that may permit selective motor protein translocation and, possibly, further regulation via

MAP-motor interactions (*43*). The organization of the outer surface of the transition zone doublet thus suggests how it could serve as a selective filter for anterograde IFT trains, by: i) directing them away from the A-microtubule and towards the B-microtubule (*53*); ii) traversing MAP9-decorated B-microtubule protofilaments B02-B06, potentially in coordination with membrane-associated transition zone complexes (*54*); and/or iii) freely walking along protofilament B01 at the AB outer junction, which is absent of MAP densities and is the site of previously described linkages between ciliary doublets and IFT trains (*16, 55, 56*). In contrast, retrograde IFT, which is regulated primarily at the ciliary tip (*57*), may tolerate less stringent selection during exit from the cilium, explaining why dynein-2 binding sites are generally more accessible on the B- (and possibly A-) microtubule.

## Acknowledgements

We thank M. Peterek and B. Qureshi from ScopeM (ETH Zürich) for data collection support. We thank M. Pilhofer and all members of the Pilhofer lab for helpful discussions. We thank F. Eisenstein for support with cryo-ET data collection. This study includes calculations performed on the Euler cluster of ETH Zürich. We acknowledge the ETH imaging platform ScopeM for instrument access.

## Author contributions

Conceptualization: B.C. and M.W.

Project administration: M.W and J.X.

Methodology: B.C., J.X., and M.W.

Resources: B.C., P.B., Y.X., J.B., A. Akhmanova, J.X., and M.W.

Investigation: B.C., J.X., A.P., E.J.G., T.V., A. Aher, E.A., and M.W.

Software: B.C., and J.X.

Formal analysis: B.C., and M.W.

Data curation: B.C., A.P., E.J.G., T.V., and M.W.

Validation: B.C., and M.W.

Visualization: B.C., and M.W.

Supervision: J.X., and M.W.

Writing - original draft: B.C., and M.W.

Writing - review and editing: M.W., B.C., and J.X.

Funding acquisition: M.W.

## Data and materials availability

The following cryo-tomograms have been deposited at the Electron Microscopy Data Bank: EMD-58112. The following cryo-ET STA maps have been deposited at the Electron Microscopy Data Bank: EMD-58085, EMD-58086, EMD-58087, EMD-58088, EMD-58089, EMD-58090 and EMD-58129. The following cryo-EM SPA maps have been deposited at the Electron Microscopy Data Bank: EMD-58091. Models have been deposited in the PDB (PDB ID: 30XP). Source data are provided with this paper. All data needed to evaluate the conclusions in the paper are present in the paper and/or the Supplementary Materials.

## Methods

### Isolation of intact cilia from bovine respiratory tract

Fresh bovine trachea was acquired from a slaughterhouse (Stadt Zürich Umwelt- und Gesundheitsschutz Veterinärdienste). The excessive tissue attached to trachea was removed and the interior of trachea was rinsed with cold PBS to remove mucus. The trachea was split into two pieces vertically to expose the respiratory epithelium. Excessive amount of cold PBS was applied to rinse the respiratory epithelium to remove remaining mucus, a nylon brush was used to gently brush the respiratory epithelium to help the removal of mucus. After the removal of mucus, remaining PBS was drained from the trachea, a T-shaped plastic Drigalski spatula was dipped into lysis buffer (20 mM HEPES-KOH pH 7.4, 25 mM KCl, 1mM EDTA, 0.25 mM Sucrose, 0.05 % Triton X-100) and used to rub against the respiratory epithelium gently to release the apical membrane of multiciliated cells. After applying lysis buffer to all areas, the respiratory epithelium was rinsed with lysis buffer, the rinse solution was collected and transferred to 50 mL centrifuge tubes. The apical membrane was pelleted at 1000 x g for 10 minutes at 4 °C, the apical membrane pellet was resuspended in 1 mL of lysis buffer and transferred to 1.5 mL centrifuge tube. To remove red blood cell contamination, the apical membrane was repeatedly resuspended in 1 mL of lysis buffer by pipetting and pelleted by centrifugation until no red blood cell contamination is visible. The apical membrane free of red blood cell was sheared 10 times using a 1 mL syringe with 26G needle. After shearing, the mixture was centrifuged at 1000 x g for 2 minutes at 4 °C to collect the supernatant, the supernatant was transferred to a new 1.5 centrifuge tube and pelted at 17,000 x g for 5 minutes at 4 °C. The pellet was resuspended in vitrification buffer (20 mM HEPES-KOH pH 7.4, 25 mM KCl, 1 mM EDTA) and inspected using phase-contrast microscopy before vitrification.

### Purification of recombinant CAPSL

A plasmid containing 6xHis-SUMO-TEV-CAPSL in the pET-24d vector was generated via gene synthesis (Genewiz). The canonical CAPSL protein isoform with UniProt accession number Q8WWF8 was used. To purify CAPSL, the plasmid was transformed into BL21(DE3) pRIL E. coli cells (Stratagene). Proteins were expressed in bacteria grown in 0.5 L of LB broth after reaching an OD600 of 0.6 in the presence of 0.5 mM IPTG at 18°C and for 16 hr. Cells were harvested by centrifugation at 4000 g at 4°C for 10 min. Harvested cell pellets were resuspended in 15 ml ice-cold lysis buffer (50 mM HEPES, pH 7.5, 150 mM NaCl, 20 mM imidazole, 2 mM 2-mercaptoethanol and 5% glycerol) and lysed by passing through an Emulsiflex C5 microfuildizer (Avestin) 3 times at >10,000 psi. The lysate was centrifuged at 40,000 g in a Type 70 Ti rotor (Beckman) for 1 hr at 4°C. The supernatant was incubated with 0.5 mL of Ni-NTA agarose (Qiagen) pre-equilibrated with lysis buffer via nutation at 4°C for 2 hr. The resin was then washed with 60 bed volumes of lysis buffer and protein was eluted with 8 bed-volumes of elution buffer (50 mM HEPES, pH 7.5, 150 mM NaCl, 300 mM imidazole, 2 mM 2-mercaptoethanol and 5% glycerol). Total eluate was mixed with 100 µg 6xHis-TEV protease produced in-house and dialyzed with Spectra/Por™ 3 RC Dialysis Membrane Tubing 3500 Dalton MWCO against 400 ml lysis buffer overnight at 4°C. Dialyzed eluate was then run through previously used Ni-NTA agarose beads to remove cleaved tag and TEV protease, and the flow though was gel filtered over a Superdex™ 75 Increase 10/300 GL column (Cytiva) pre-equilibrated in gel filtration buffer (20 mM HEPES, 150 mM NaCl, 2 mM 2-mercaptoethanol and 5% glycerol). Fractions were analyzed by SDS-PAGE followed by Coomassie staining. Peak fractions were polled, aliquoted, snap-frozen and stored at −80°C.

### Grid preparation

For intact cilia isolated from bovine respiratory tract, 4 μl of cilia suspsension was mixed with 1 μl of 10-nm bovine serum albumin (BSA) – coated colloidal gold particles (Cytodiagnostics), 4 μl of mixed sample was applied onto negatively glow discharged EM grids (R3.5/1 Cu 200 mesh, Quantifoil) coated with a 2-nm-thick continuous carbon layer in a Vitrobot Mark IV (Thermo Fisher Scientific). After 30-s incubation time, grids were blotted for 6 to 8 seconds at 8 °C and plunge frozen in 37% ethane-propane mixture.

Alternatively, cilia suspension was diluted 6 times first, 4 μl of the diluted suspension was mixed with 1 μl of 10-nm gold particles, 4 μl of the mixture was applied onto negatively glow discharded EM grids (R3.5/1 Cu 200 mesh, Quantifoil) without carbon layer in a EM GP2 Automatic Plunge Freezer (Leica). After 30-s incubation, grids were blotted for 12 seconds at 8 °C and plunge frozen in 37% ethane-propane mixture.

To generate paclitaxel-stabilized microtubules, purified porcine brain tubulin was first diluted to 20 µM in BRB80 buffer (80 mM PIPES, 1 mM EGTA, 1 mM MgCl_2_ pH 6.8) supplemented with 2 mM GTP and 10% glycerol. Mixture was incubated for 5 minutes on ice and spun in a TLA.100 rotor at 90,000 rpm for 10 minutes at 4°C. Supernatant was incubated in a 37°C water bath for 2 minutes. Then, at 37°C, 5 µl of 1 µM paclitaxel (Sigma-Aldrich®) in DMSO were added, followed by an incubation of 10 minutes, addition of 5.5 µl of10 µM paclitaxel in BRB80, 10 minute incubation, addition of 6 µl 100 µM paclitaxel in BRB80 and a final 15 minute incubation. Taxol-stabilized microtubules were spun in a TLA.100 rotor at 90,000 rpm for 10 minutes at 30°C and resuspended in 100 µl electron microscopy paclitaxel buffer (BRB80 supplemented with 1 mM DTT, 0.05% NP-40 (Roche) and 10 µM paclitaxel). Tubulin concentration was determined by nanodrop.

To generate cryo-EM grid specimens containing CAPSL-decorated, paclitaxel-stabilized microtubules, freshly thawed CAPSL was diluted in electron microscopy paclitaxel buffer and mixed with freshly prepared paclitaxel-stabilized microtubules, to a final tubulin concentration of 0.2 mg/ml and CAPSL concentration of 19.35 µM. Mixture was incubated at 37°C for 30 minutes. 3.5 µl of the solution was pipetted onto a glow-discharged EM grid (C-Flat, Cu 300 mesh, thick) in Vitrobot Mark IV (Thermo Fisher Scientific), after 30 s of incubation at 100% humidity and 22°C. Grids were blotted for 3 s with a blot force of −1, plunge-frozen in 37% ethane-propane mixture and stored in liquid nitrogen until data collection.

### Cryo-ET data collection

For the 1,201 tomograms in dataset 1, SerialEM (*60*) and SPACETomo (*61, 62*) were used to collect data on a Titan Krios transmission electron microscope G4 (Thermo Fisher Scientific) operating at 300 kV and equipped with a K3 direct electron detector and a BioContinuum energy filter (Gatan). The targets were identified in the grid square montages at low magnification in SPACETomo target selection GUI, tilt series were collected at a nominal magnification of 64kx (1.384 Å per pixel) using a dose-symmetric scheme with an angular range of +60° to −60° and an angular step of 3°, a total does of ∼160 e^−^/Å^2^, and at a defocus ranging from −2 to −4 μm.

For the 126 tomograms in dataset 2, TOMO5 (Thermo Fisher Scientific) was used to collect data on a Titan Krios transmission electron microscope G3i (Thermo Fisher Scientific) operating at 300 kV and equipped with a K3 direct electron detector and a BioQuantum energy filter (Gatan). The targets were identified in search maps in TOMO5 GUI, tilt series were collected at a nominal magnification of 64kx (1.34 Å per pixel) using a dose-symmetric scheme with an angular range of +60° to −60° and an angular step of 3°, a total does of ∼160 e^−^/Å^2^, and at a defocus ranging from −1 to −3 μm.

### Sub-tomogram averaging

All tilt-series were created using alignframes in IMOD v5.1 (*63*) with collected frames and mdoc files. Tilt-series alignment was performed with Etomo (*64*) in IMOD v5.1 in batch mode using 10 nm gold particles as fiducial, and the alignment results were imported into WarpTools (*65*).

Motion correction CTF estimation of frames were performed in WarpTools (*65*). Tomograms were reconstructed at bin8 using the alignment result imported from IMOD. Reconstructed tomograms were denoised using CryoCARE v11 (*66, 67*).

1327 Denoised tomograms were imported into Dynamo (*68, 69*), the microtubule triplets in basal body and transition zone microtubule doublets were manually picked as filament models using dtmslice GUI and extracted with z step of 8 nm. 680,495 subtomograms were picked and the resulting table file from Dynamo (*68, 69*) were transformed to Relion3 (*70*) format star file, the TiltPrior/PsiPrior were calculated based on the geometrical position of neighboring particle in each filament, the HelicalTrackLength was scaled using the pixel size to HelicalTrachLengthAngst.

Particles were extracted in WarpTools at bin8 as 3D subtomograms and imported into Relion4 (*71*). Using the microtubule triplet as initial reference, the extracted 3D subtomograms were subjected to 3D classification with only 1 class, the angular search was set to search Rot globally and Tilt/Psi locally around the TiltPrior/PsiPrior for initial alignment. The initially aligned subtomograms were grouped into sub-zones based on the HelicalTrackLength of each subtomogram, every 50 nm of HelicalTrackLength is considered as a sub-zone. The 3D subtomgorams were splitted into sub-zones, relion_reconstruct was used to create reconstructions of each sub-zone. 15 sub-zones were created, and 15 classes were reconstructed to be used as references for subsequent supervised 3D classification. The roughly aligned particles were subjected to 3D classification (without alignment), the 15 classes of sub-zones were used as references to classify the subtomograms into 15 classes. 93,174 subtomograms from classes with clear transition zone doublet features were selected and refined using 3D refinement.

The 3D refinement job was imported to MCore for high resolution refinement, one specie was created with a pixel size of 4 Å and subjected to multiple rounds of MCore (*72*) refinement to correct imagewarp, volumewarp, ctf, stageangle and anisotropic magnification. The MCore refinments updated the .xml files for each tomogram, and the 3D subtomograms were exported in WarpTools at bin8 after updating the .xml files, the new 3D subtomograms were subject to 3D classification (withtout alignment) in Relion4. Selected particles were subject to 3D refinement and repeated MCore refinement as above.

After the final round of 3D classification and 3D refinement in Relion4, 51,069 subtomograms were selected and the 3D refinement job was imported to MCore for high resolution refinement, one specie was created with a pixel of 3 Å and performed refinements including imagewarp, volumewarp, ctf, cs, stageangle and anisotropic magnification. The refinement result was used to create species at pixel size of 2 Å and 1.384 Å for further refinement in MCore. After refinement for species with pixel size of 1.384 Å, the resolve_items and resolve_frames in EstimateWeights (*65*) were performed and an additional round of MCore refinement was performed.

For local refinement, for species at pixel size of 1.384 Å were created and centered on A03, A08, B02 and B06 protofilament using shift_species in MTools (*72*). Additional rounds of MCore refinements were performed to improve the local resolution. Refinement results were postprocessed using Relion4 to calculate the resolution and gold-strandard FSC, postprocessed maps were sharpened using EMReady v2.3 (*59*) to enhance the features.

### Cryo-EM data collection

Cryo-EM single-particle analysis dataset was collected using EPU (Thermo Fisher Scientific) on a Titan Krios transmission electron microscope G3i (Thermo Fisher Scientific) operating at 300 kV and equipped with a K3 direct electron detector and a BioQuantum energy filter (Gatan). Automated collection was performed at a nominal magnification of 64kx (1.34 Å per pixel). The total electron dose applied was 42 e^−^/Å^2^, at a defocus ranging from −1.0 to −3.0 μm.

### Cryo-EM data processing

Motion correction of 8,036 raw movies was performed using MotionCor2 (*73*) in Relion5 (*74, 75*). Motion-corrected micrographs were imported into CryoSPARC v5.0 (*75, 76*), PatchCTF and FilamentTracer (reference free) were performed to extract 1.03M microtubule particles in filament mode at bin4. Extracted particles at bin4 were cleaned with multiple rounds of 2D classification, selected particles were classified by Heterogenous Refinement using microtubules composed of 11 to 16 protofilaments as initial reference (*77*). 165k particles with 13 protofilaments were selected and exported to Relion5 using pyem (*78*) to perform 3D classification (without alignment), classes with clear CAPSL-decoration containing 36k particles were selected. The reconstruction result of the selected 3D class was imported into CryoSPARC v5.0 as the reference for FilamentTracer.

A second round of FilamentTracer was performed using the CAPSL-decorated microtubule reconstruction as reference. Extracted particles at bin4 were cleaned with multiple rounds of 2D classification, 464k particles were selected.

The selected 464k particles were classified by Heterogenous refinement using microtubule composed of 13 and 14 protofilaments. Particles with 13 protofilaments were selected.

To better classify the CAPSL density and identify the seam of 13 protofilament microtubule, the selected particles were used for 2D multicurve fitting and microtubule-signal subtraction with a scale_factor of 0.3 to weaken the microtubule lattice signal (*79*). Subtracted micrographs were imported back to CryoSPARC v5.0 and performed PatchCTF, the information of the selected particles from the Heterogeneous Refinement job was exported in star file format using pyem and re-imported into CryoSPARC v5.0 to establish links with the subtracted micrographs. The particles were re-extracted at bin4 from the subtracted micrographs and classified with 2D classification. 395k particles were selected for heterogenous Refinement using CAPSL-decorated 13-protofilament microtubule and naked 13 protofilament microtubule as reference. Selected 77k particles were reextracted at bin2, after removing duplicated particles, 58,610 particles at bin2 were exported to Relion5 using pyem.

In Relion5, 3D classification was performed with a mask focusing on the seam region to select classes with obvious CAPSL mismatch at the seam. Selected 35,389 particles were re-imported into CryoSPARC v5.0. The particles were reextracted at bin1 and refined using Helical Refinement (without helical symmetry) and Local Refinement. The Local Refinment result were postprocessed in Relion5 and sharpened using EMReady v2.3 to enhance the features of the map.

### Protein identification

The CAPSL, ENKD1 and FAM161B protein were identified using CryoAtom. The final local refinement results from MCore were postprocessed using Relion4 and used as input for CryoAtom. All reviewed Uniprot entries of bovine protein sequences were downloaded from Uniprot as a single FASTA file as input for CryoAtom. CryoAtom were ran directly on the maps with the bovine protein sequence database provided. CAPSL, ENKD1 and FAM161B were assigned to the maps by CryoAtom, structural predictions of tubulin dimer with CAPSL, ENKD1 and FAM161B were done using AlphaFold3 to assist model building.

The LC8 protein was identified using DomainSeeker. All the Uniprot accession numbers of reviewed human protein sequences were downloaded from Uniprot, which is used to pull AlphaFold2 results including the pdb and pae files from the AlphaFold Protein Structure Database (*26*). The pdb files were parsed into domains using corresponding pae files. The globular domains of MAPs from the transition zone doublet density were cropped in ChimeraX and used as input for domain fitting in DomainSeeker. The domain fitting results were scored using DomainSeeker, which assigned LC8 to the globular density on A06 protofilament which is formed by LC8 dimer.

The DZANK1 protein assignment was inspired by the interaction between LC8 (DYNLL1/DYNLL2) and DZANK1 (*80*). AlphaFold3 prediction of DZANK1 and LC8 dimer suggestion that DZANK1 forms a complex with LC8 dimer, which contains an α-helix that matches the density of MAPs in between A06 and A07 protofilament. AlphaFold3 prediction of DZANK1 dimer also suggests the globular density on A07 also fits well with DZANK1 dimer.

The ECT2L protein assignment was based on DZANK1 assignment. The DZANK1 interacting proteins were pulled from the Human Predictome database (*81*). As LC8 and DZANK1 forms dimers, the monomeric and homo-dimeric structures of DZANK1-interacting proteins were predicted. All predicted structure were parsed using the pae files from monomeric predictions. The parsed domains were fitted into the globular domain on B06 protofilament. ECT2L dimer was fitted into the B06 globular density with multiple α-helices.

The RPGRIP1L and SPATA7 were assigned using AlphaFold3, by prediction the multimer structure of tubulin dimer with proteins that were reported to be related to the transition zone or the connecting cilium in photoreceptor (*7, 82–85*). The prediction containing tubulin dimer with RPGRIP1L and SPATA7 fitted well into the density of MAPs on A09-A10 protofilament and B02 protofilament.

The CEP41, CFAP20 and MAP9 protein assignment were based on homology with published structures (*15, 18, 43*).

### Immunofluorescence cell staining of human nasal epithelial cells (HNECs)

Nasal brushes were collected from healthy human patients with explicit broad consent for biobanking in the UMC Utrecht Airbank (approved by ‘Toetsingscommissie biobanking (TcBio)’ protocol ID 16-586). Experiments with these cells were performed under the approved protocol Tcbio ID 22-079. Culture and maintenance of human nasal epithelial cells were described previously, using materials of wild-type and immortalized Human Nasal Epithelial Cell (HNECs) donor HNEC3 (*23, 35*). In brief, HNECs were washed with PBS for 5 minutes prior to fixation. The samples in this study were fixed with either ice-cold MeOH at −20°C for 20 min or 4% paraformaldehyde and 4% sucrose in MRB80 (80 mM PIPES pH 6.8, 1 mM EGTA and 4 mM MgCl2) for 20 min at 37°C. Post-fixation samples were subjected to 3 PBS washes of at least 10 min each. Washed cells were blocked with 3% BSA in PBS for at least 1 h at room temperature. Filter pieces were incubated in a 0.5 ml Eppendorf tube with 30 μl primary antibody diluted with 3% BSA in PBS at 4°C overnight with constant agitation. Then, cells were washed by 3 PBS washes of 10 min and incubated in secondary antibody as described for primary antibody incubation for at least 3 h at room temperature. After secondary antibody incubation, cells were subjected to 3 PBS washes of 10 min, cells were dehydrated in 70% EtOH followed by 100% EtOH and mounted in ProLong Diamond or Vectashield using 13 mm thickness 1.5 coverslips.

The primary antibodies used for immunofluorescence cell staining of target proteins are rabbit antibodies: CAPSL (HPA046811), FAM161B (HPA019125), RPGRIP1L (HPA040530) and ENKD1 (HPA041478). The secondary antibody used is Goat anti-Rabbit IgG (H+L) Highly Cross-Adsorbed Secondary Antibody, Alexa Fluor™ 594 - A-11037.

The primary antibody used for immunofluorescence cell staining of tyrosinated tubulin was Rat Alpha Tubulin Monoclonal Antibody (YL1/2) - MA1-80017. The secondary antibody used was Goat anti-Rat IgG (H+L) Cross-Adsorbed Secondary Antibody, Alexa Fluor™ 488 - A-11006.

### Expansion microscopy

Methods for the culture and maintenance of human nasal epithelial cells were described previously (*23, 86*). In brief, the fixation and staining methods are the same as the Immunofluorescence cell staining. After the removal of the secondary antibody solution, MeOH fixed cells were post-fixed with 4% paraformaldehyde in PBS for 10 min at room temperature (post-fixation was not required for PFA-fixed cells) and incubated in 0.1 mg/ml acryloyl X-SE and 0.01% Triton X-100 in PBS overnight at room temperature with constant agitation. After 2 PBS washes of 15 min, filter pieces were placed onto a parafilm-covered glass slide with cells facing up, covered by a plastic Transwell scaffold and sealed with grease. To this gelation chamber, 75 μl gelation solution (1% w/v sodium acrylate, 14.4% w/v acrylamide, 0.009% N, N’-methylenebisacrylamide, 1x PBS, 0.1% Triton X-100, 0.15% tetramethylethylenediamine (TEMED) and ammonium persulfate (APS) in MiliQ water) was added and allowed to polymerize for 1 h at 37°C. The gel was carefully removed from the Transwell scaffold and the gel section containing the filter piece was trimmed using a scalpel. Then, the gel was transferred to a 12-well plate for homogenization in digestion solution (0.5% Triton X-100, 0.8 M guanidine-HCl, 9 U/ml Proteinase K in TAE (40 mM Tris, 20 mM aceticacid, 1 mM EDTA in MiliQ water) for 4 h at 37°C. After homogenization, the gel was thoroughly washed 3 times 15 min with PBS and in case a total protein stain was used, transferred to a 24-well plate and incubated in 20 μg/ml ATTO-NHS-643 in PBS for 1.5 h at room temperature with constant agitation. During the last 15 min of incubation, 5 μg/ml DAPI was added. To expand the sample, gels were transferred to a 15 cm petri dish with MiliQ water and incubated at room temperature. After 30 min, the MiliQ water was refreshed, and the sample was expanded overnight. Prior to imaging, the gels were trimmed and mounted onto a plasma-cleaned, poly-L-lysine treated cover glass.

The primary antibodies used for staining the target proteins in expansion microscopy were the rabbit antibodies: MAP9 (26078-1-AP) and SPATA7 (HPA038082). The secondary antibody used was Goat anti-Rabbit IgG (H+L) Highly Cross-Adsorbed Secondary Antibody, Alexa Fluor™ 488 - A-11034.

### Image acquisition

Confocal images for non-expanded immunofluorescence samples were acquired using a Zeiss LSM700 microscope equipped with 405 nm, 488 nm, 555 nm and 633 nm lasers and Plan-Apochromat 63x/1.40 Oil DIC objective.

Expanded samples were imaged using a Leica TCS SP8 STED 3X microscope with continuous 405 nm and pulsed (80 MHz) white light lasers, PMT and HyD detectors and spectroscopic detection with a HC PL APO 86x/1.20W motCORR STED (Leica 15506333) water objective.

### TIRF-based microtubule nucleation assays

The microscope setup used was a Zeiss Elyra 7 microscope equipped with an α Plan-APOCHROMAT 63x/1.46 Oil Corr TIRF objective with a numerical aperture of 1.46, operated under immersion oil (Zeiss), 2x Hamamatsu ORCA-Fusion BT (QE around 95 %), and 4 lasers generating excitation at 405 nm, 488 nm, 561 nm and 640 nm. Image acquisition was controlled using Zen Blue 3.1 (Zeiss).

Glass slides (Premiere 8201) and glass coverslips (Corning, 22 × 22 mm, 2845–22) were rinsed in acetone, followed by water, sonicated for 20 min in 50% methanol, rinsed with water, sonicated for 20 minutes in 0.5 M KOH and rinsed with water before drying using filtered compressed air. Then, the clean, dry glass was subjected to plasma cleaning for 10 minutes at high intensity in a Harrick Plasma PDC-32G-2 plasma cleaner connected to an ICME M71B4 vacuum pump.

For experiments in fig. S10, the channel was rinsed with 1X BRB80 buffer followed by the application of native γ-TuRC peak fractions (purified as in (*87*)) diluted 5-fold in 1X assay buffer (1X BRB80(80 mM PIPES, 1mM EGTA, 1mM MgCl_2_, pH 6.80 with KOH) - 1 mM TCEP, 50 mM KCl, 0.15% [weight/volume] methylcellulose, 0.2 mg/mL κ-casein and 1 mM GTP). After 5 minutes of incubation, non-adherent molecules were washed away by 1X assay buffer. A 15 μM solution of fluorescently labeled red tubulin in 1x assay buffer containing an oxygen scavenger system (0.035 mg/mL catalase, 0.2 mg/mL glucose oxidase, 2.5 mM glucose, and 10 mM DTT) introduced to the flow cell with or without 1 μM Calcyphosin like (CAPSL). Immediately afterwards, the flow cell was sealed with VALAP (1:1:1 vaseline, linoleum, paraffin wax) and placed on the preheated (37°C) microscope stage. Each sample was measured for 20 minutes in 5 s intervals with excitation at 561 nm (10 ms exposure).

---

**Figure S1.**
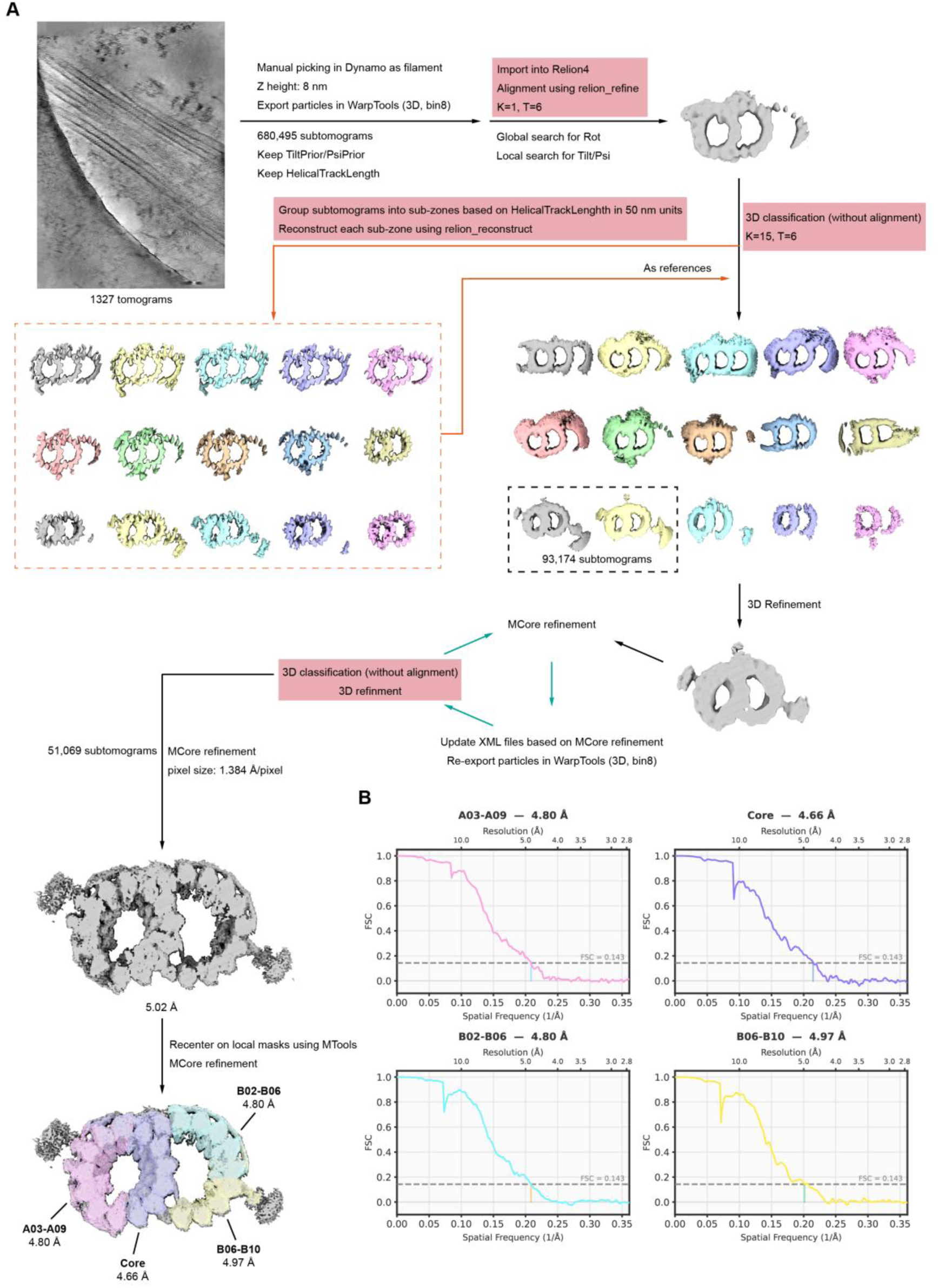
STA processing of the transition zone doublet from intact bovine cilia. **A.** Top left: Slice through a representative cryo-tomogram of the transition zone in an isolated bovine trachea cilium (10 nm projection), showing multiple microtubule doublet structures. The rest of this panel presents a workflow for the cryo-ET STA processing scheme, which includes steps in both RELION-4 (*71*) (steps coloured in red) and Warp/M (*65, 72*) (see Materials and Methods for processing details). **B.** Gold-standard Fourier Shell Correlation (FSC) plots of the local transition zone doublet reconstructions shown at the bottom left of panel **A**, centered around protofilaments A03-A09 (pink curve), the “core” (purple curve), protofilaments B02-B06 (blue curve), and protofilaments B06-B10 (yellow curve).

**Figure S2.**
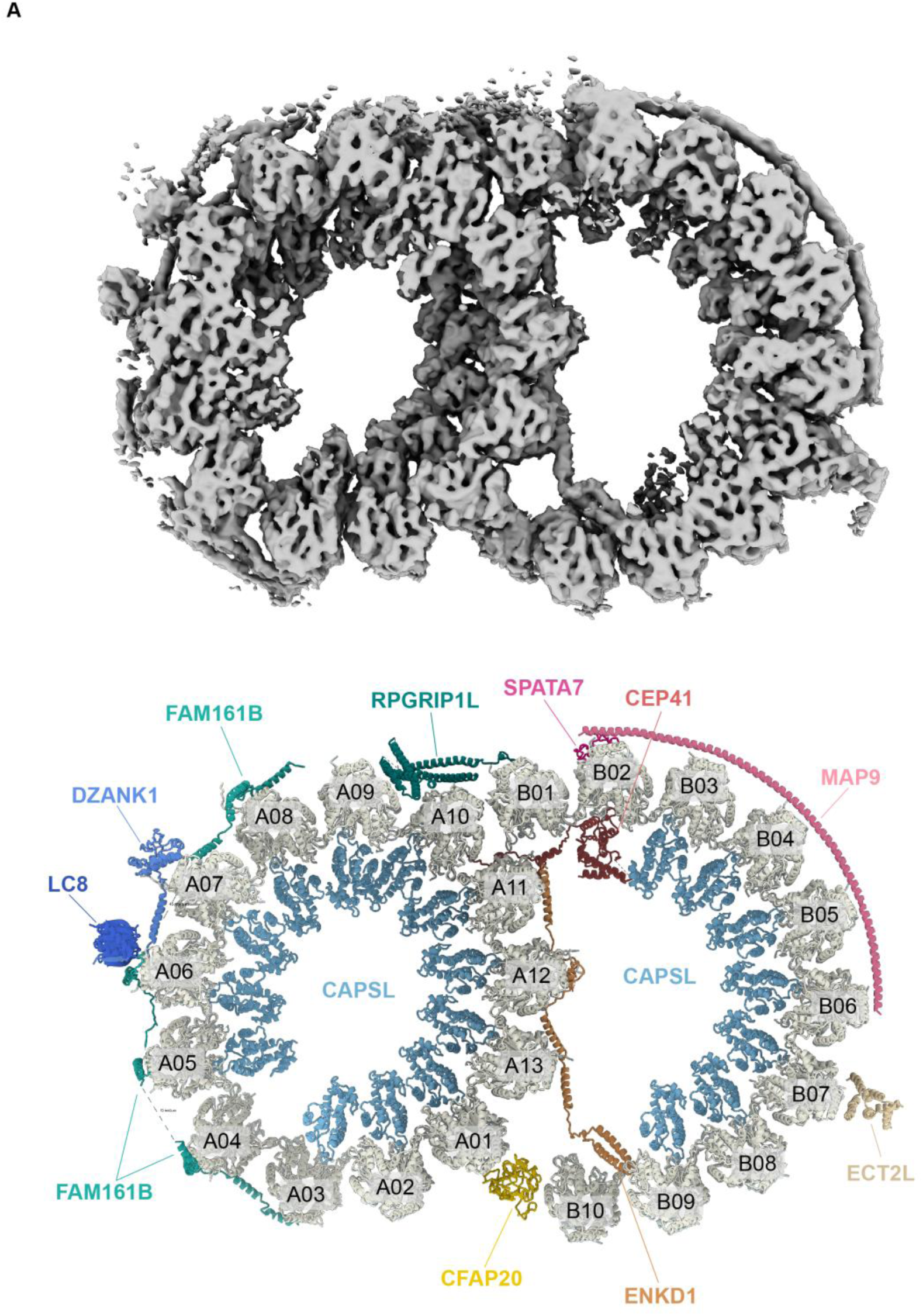
Molecular model of the transition zone doublet microtubule.

**Figure S3.**
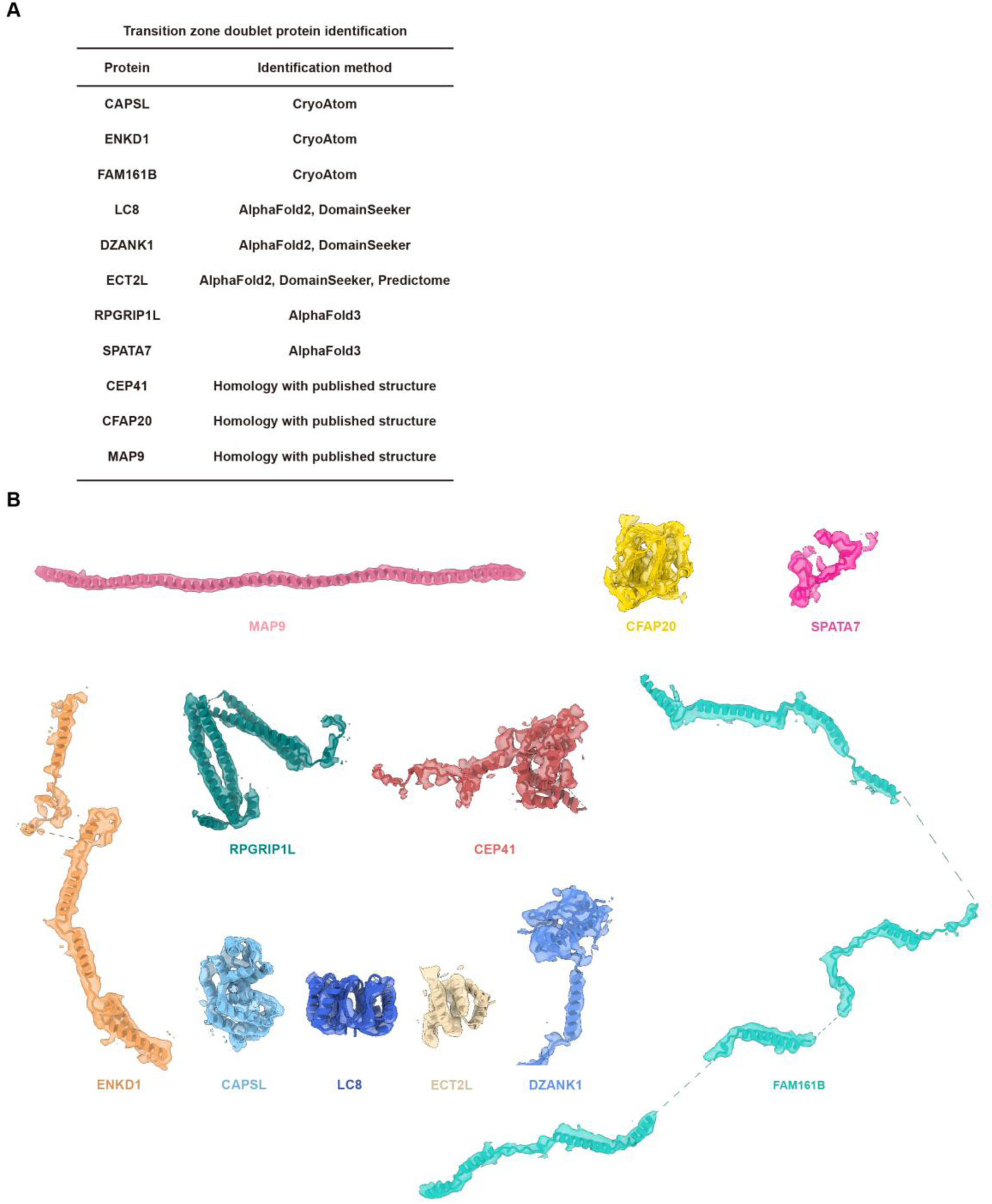
The identification of MIPs and MAPs in the transition zone doublet. **A.** A list of the identification methods used for assigning each indicated protein to the transition zone doublet. See Materials and Methods for further assignment details. **B.** Refined transition zone doublet MIP and MAP models generated in this study (cartoon representation) shown in corresponding segmented densities from the STA reconstruction (transparent surfaces). Subunits and densities are colored as in Figure 1.

**Figure S4.**
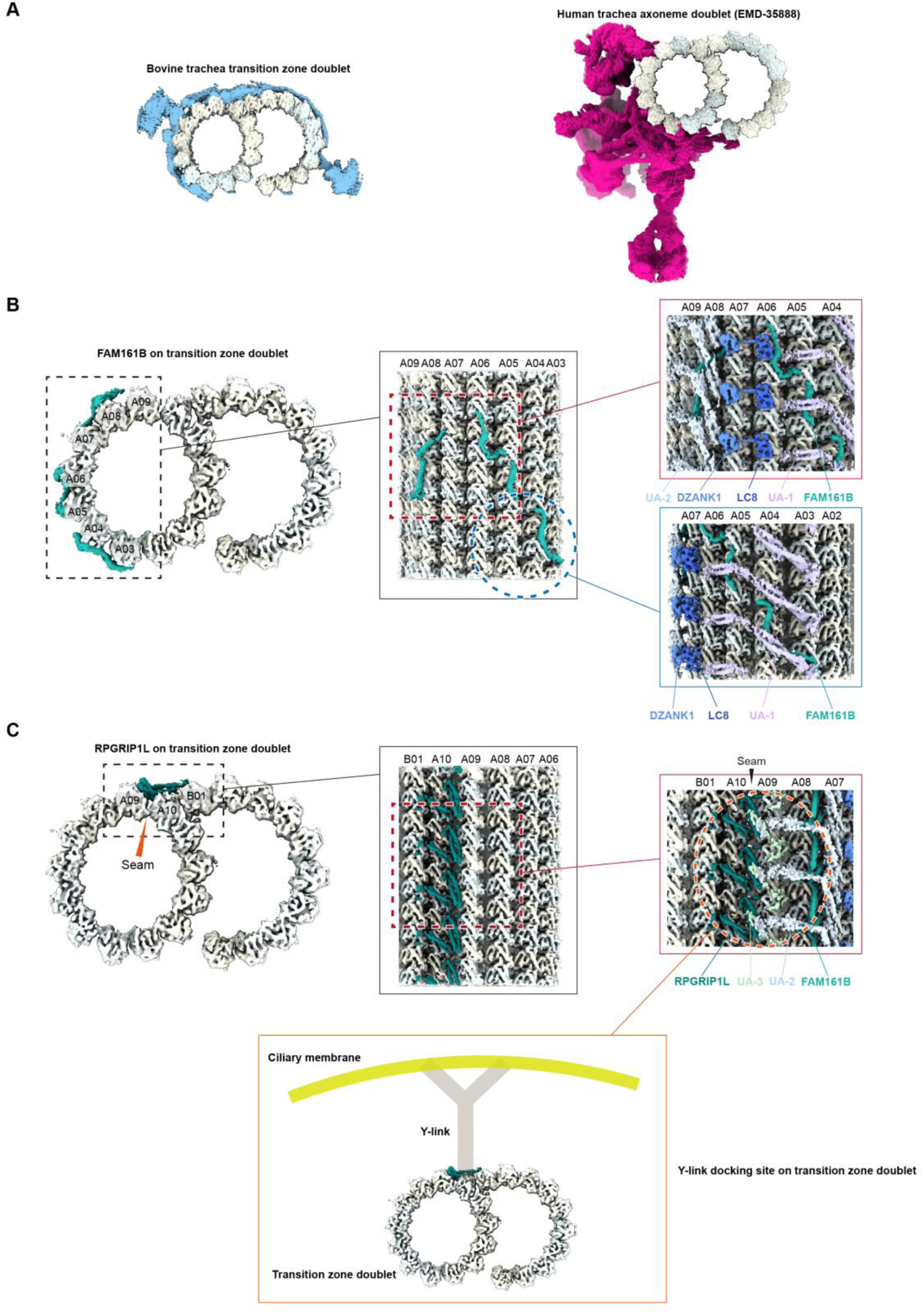
The A-microtubule MAPs in the transition zone doublet. **A.** Left: Cross section view of the transition zone doublet density map (this study) including its 8-nm repeat MAP network (blue density; tubulin heterodimers shown in ivory/azure). Right: Cross section view of the axonemal doublet density map (EMD-35888) (*88*) including its 96 nm repeat MAP network (pink density, including dyneins; tubulin heterodimers shown in ivory/azure). **B.** Left: Cross section view of the transition zone doublet density map showing only density for tubulin and FAM161B. Protofilaments A03-A09 are indicated to show the span of FAM161B’s association across the A-microtubule.Middle: Rotated view of the dashed region on the left facing the entire FAM161B density. Right: Views of the dashed regions in the middle panel without segmenting any densities to highlight the MAP network that sits atop of FAM161B near the TZ-linker (top) or towards the AB inner junction (bottom). Subunit densities are colored as in Figure 1. **C.** Left: Cross section view of the transition zone doublet density map showing only density for tubulin and the RPGRIP1L dimer. Protofilaments A09, A10 and B01 are indicated to show how RPGRIP1L spans both the A-microtubule seam (indicated) as well as the AB outer junction.Middle: Rotated view of the dashed region on the left facing the RPGRIP1L density.Right: View of the dashed region in the middle panel without segmenting any densities to highlight the MAP network adjacent to RPGRIP1L. Subunit densities are colored as in Figure 1.Bottom: Cross section view of the transition zone doublet density map showing only density for tubulin and the RPGRIP1L dimer together with a schematic of the Y-link and its attachment to the ciliary membrane to highlight how the Y-link docking site on the transition zone doublet corresponds near the RPGRIP1L density.

**Figure S5.**
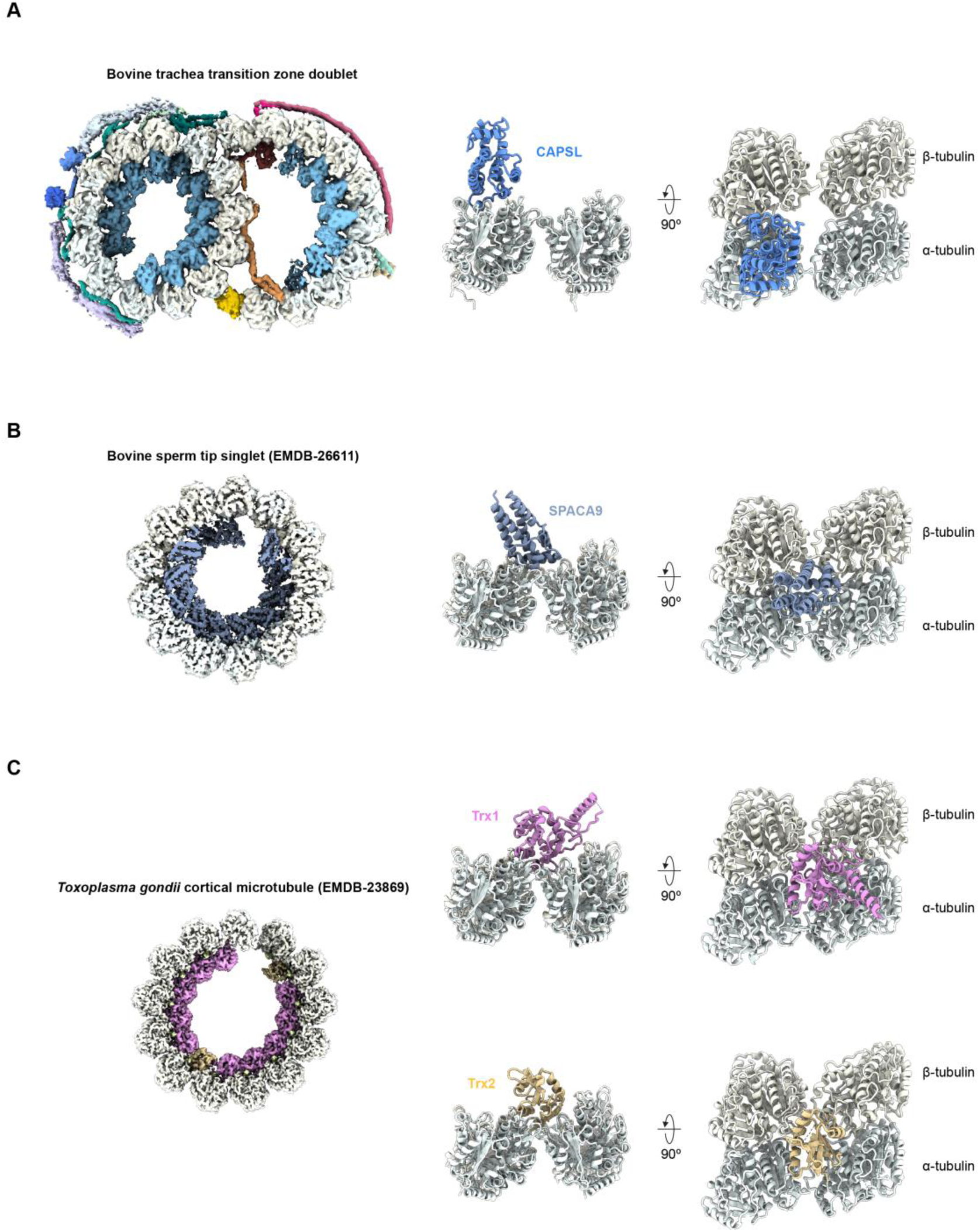
CAPSL binding mode on tubulin compared with other MIPs in bovine cilia. **A.** Left: Cross-sectional view of the STA transition zone doublet composite map. Protein densities are colored based on protein identity assignment, as in Figure 1.Right: Two views of CAPSL bound relative to two neighboring tubulin dimers, taken from the transition zone doublet model (cartoon representation; the second CAPSL is omitted for clarity). **B.** Left: Cross-sectional view of the bovine sperm tip singlet reconstruction (EMD-26611) (*31*). Tubulin density is colored in white, SPACA9 density is colored in blue. Right: Two views of SPACA9 bound relative to two neighboring tubulin dimers, taken from PDB ID: 7UN1 (cartoon representation) (*31*). **C.** Left: Cross-sectional view of the *Toxoplasma gondii* cortical microtubule reconstruction (EMD-23869) (*34*). Tubulin density is colored in white, Trx1 density is colored in pink, and Trx2 density is colored in beige. Right: Two views of Trx1 (top) or Trx2 (bottom) bound relative to two neighboring tubulin dimers, taken from PDB ID: 7MIZ (cartoon representation) (*34*).

**Figure S6.**
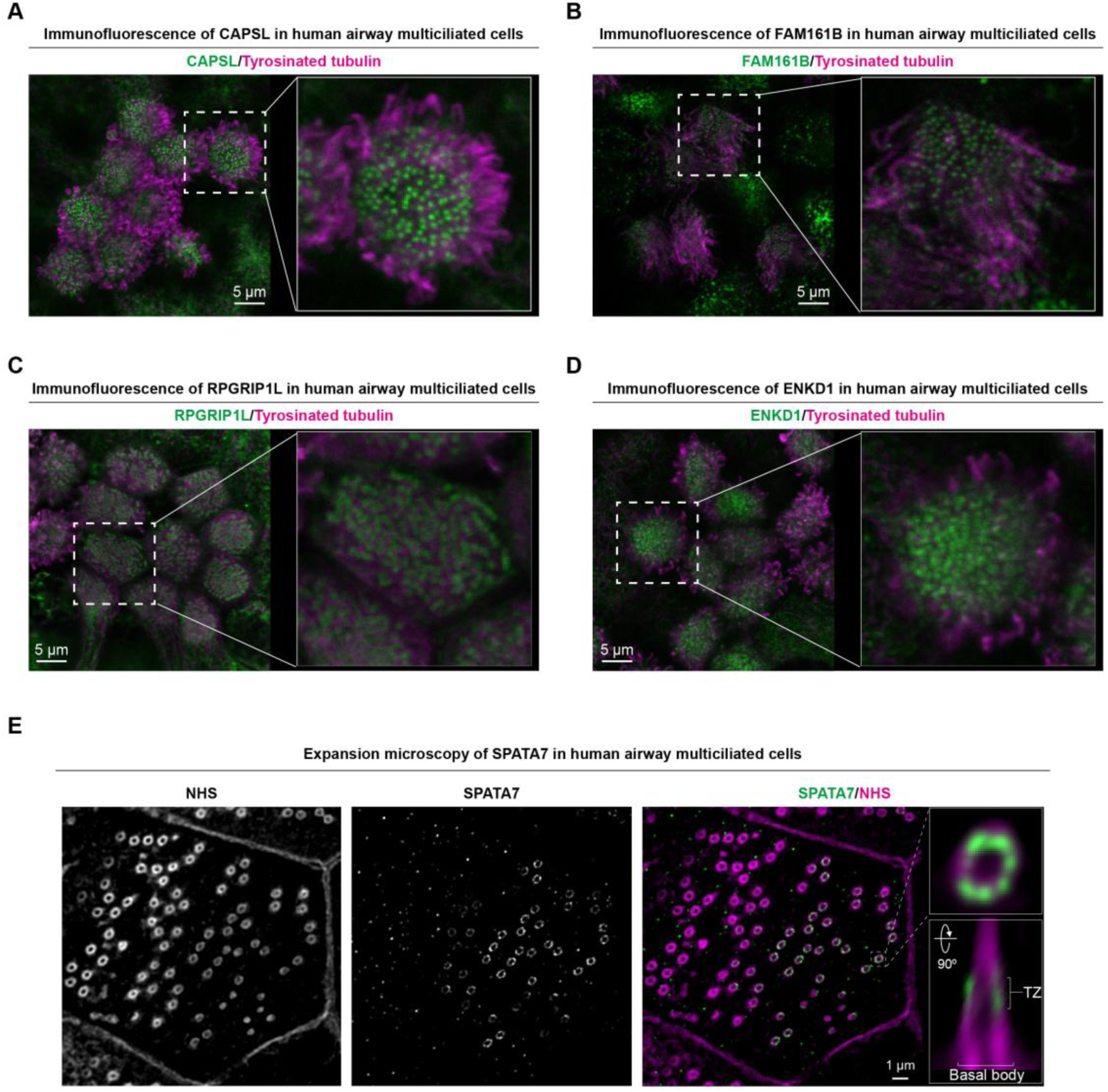
Fluorescence microscopy analysis of identified transition zone doublet MIPs and MAPs. **A.** Immunofluorescence microscopy images of HNECs stained for CAPSL. **B.** Immunofluorescence microscopy images of HNECs stained for FAM161B. **C.** Immunofluorescence microscopy images of HNECs stained for RPGRIP1L. **D.** Immunofluorescence microscopy images of HNECs stained for ENKD1. **E.** TREx (*86*) images of HNECs stained for whole protein via fluorescent NHS (left) or SPATA7 (middle). A merged image is shown on the right, with insets showing orthogonal cross sections across or along the transition zone. The location of the basal body is also indicated.

**Figure S7.**
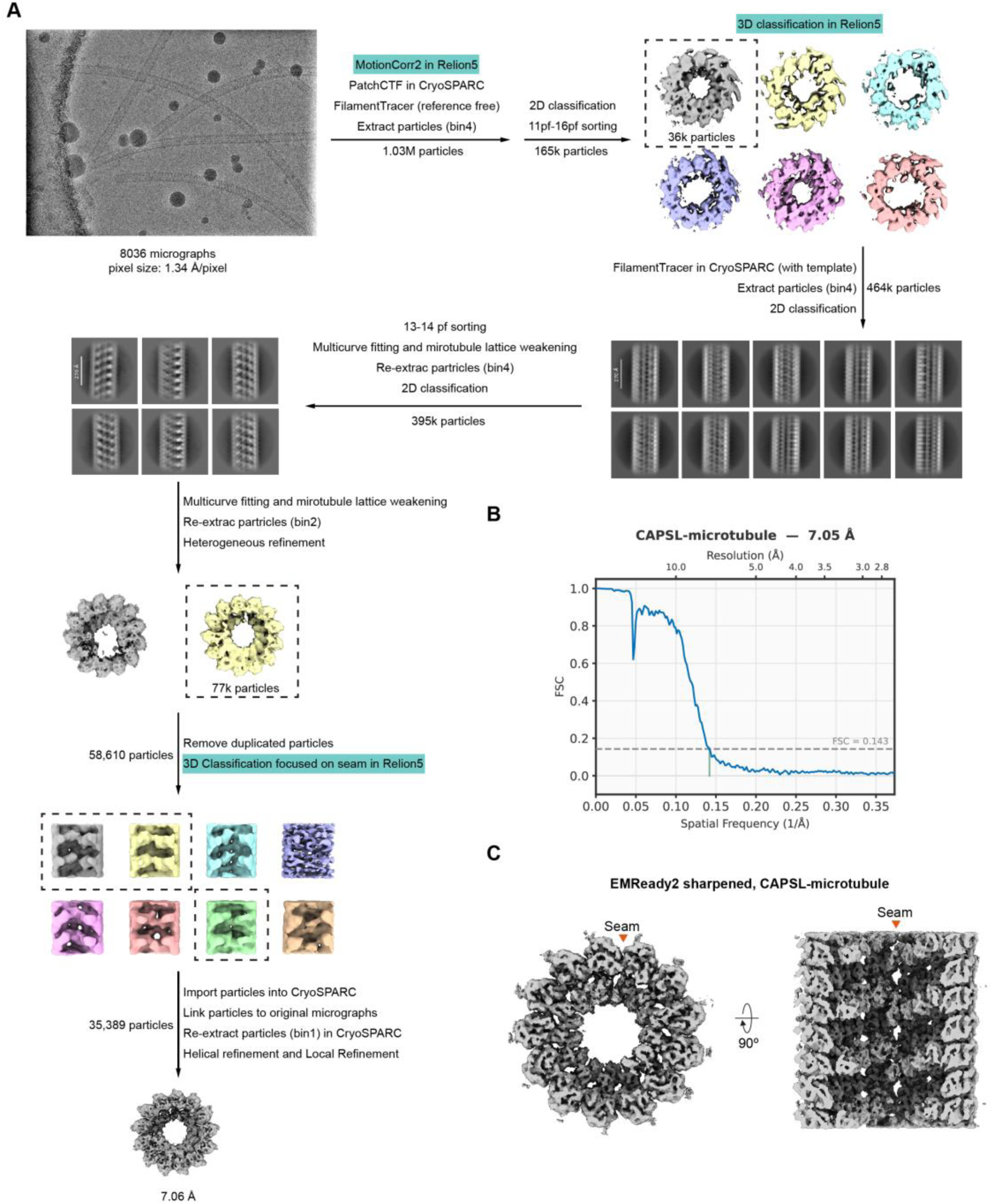
SPA processing of human CAPSL-decorated, paclitaxel-stabilized microtubules. **A.** Top left: Representative cryo-EM micrograph of porcine brain paclitaxel-stabilized microtubules decorated with purified recombinant human CAPSL. The rest of this panel presents a workflow for the SPA processing scheme, which includes steps in both CryoSPARC v5.1 and RELION-5.1 (steps colored in green) (see Materials and Methods for processing details). **B.** Gold-standard Fourier Shell Correlation (FSC) plot of the CAPSL-microtubule reconstruction. **C.** Two orthogonal cross-sectional views of the 13-protofilament CAPSL-microtubule reconstruction with the location of the seam indicated. Map was sharpened using EMReady2 (*59*).

**Figure S8.**
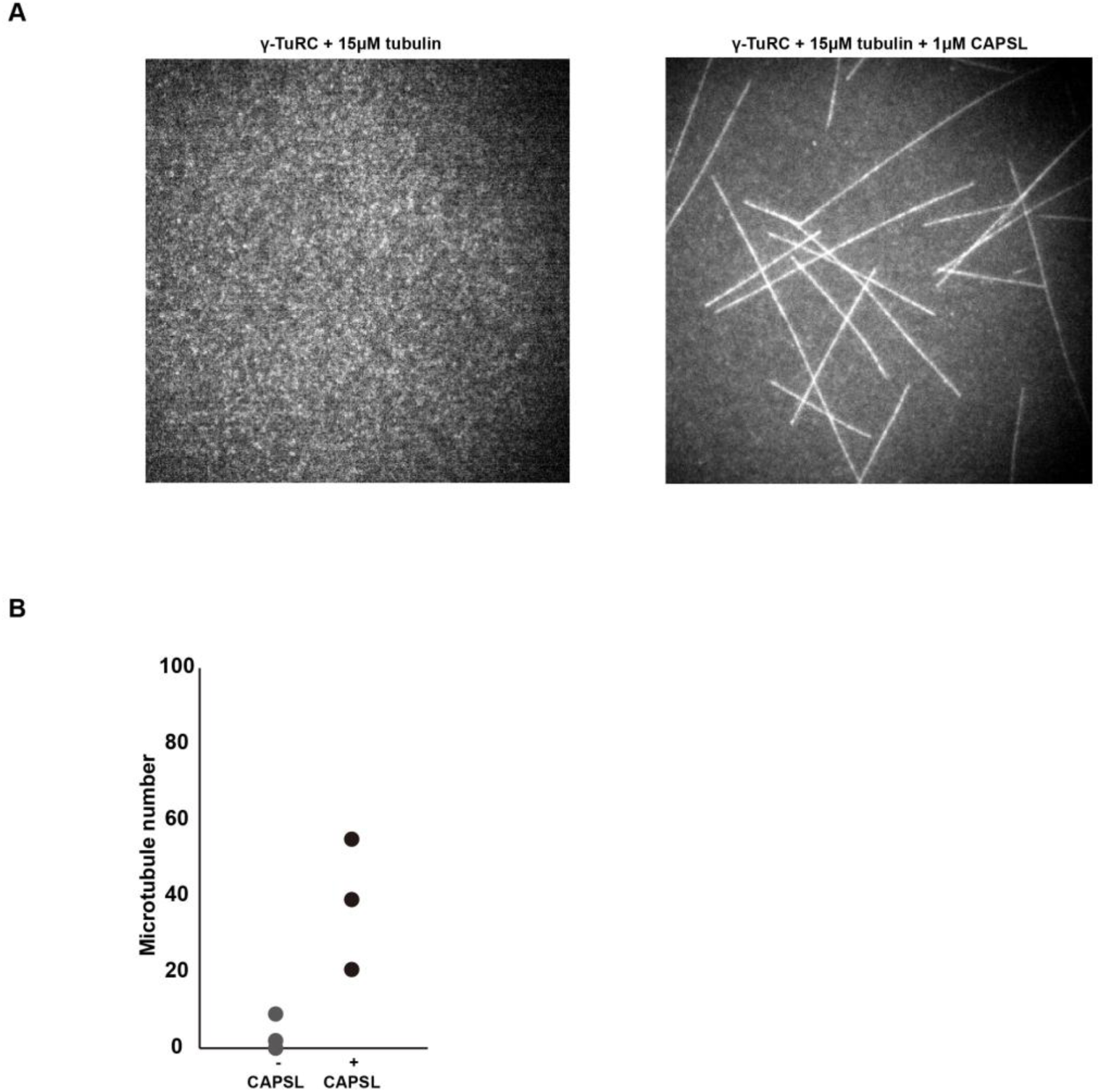
Effect of CAPSL on microtubule assembly *in vitro* visualized using TIRF microscopy. **A.** Fields of view of microtubule assembly after ∼20 minutes at 15 μM fluorescently labeled tubulin nucleated by surface-adsorbed, non-fluorescent human γ-tubulin ring complexes in the absence (left) or presence (right) of 1 μM CAPSL. **B.** Plot of the number of microtubules after ∼20 minutes in the experiments in panel **A**. Three independent experiments were conducted for each condition on two separate days.

**Figure S9.**
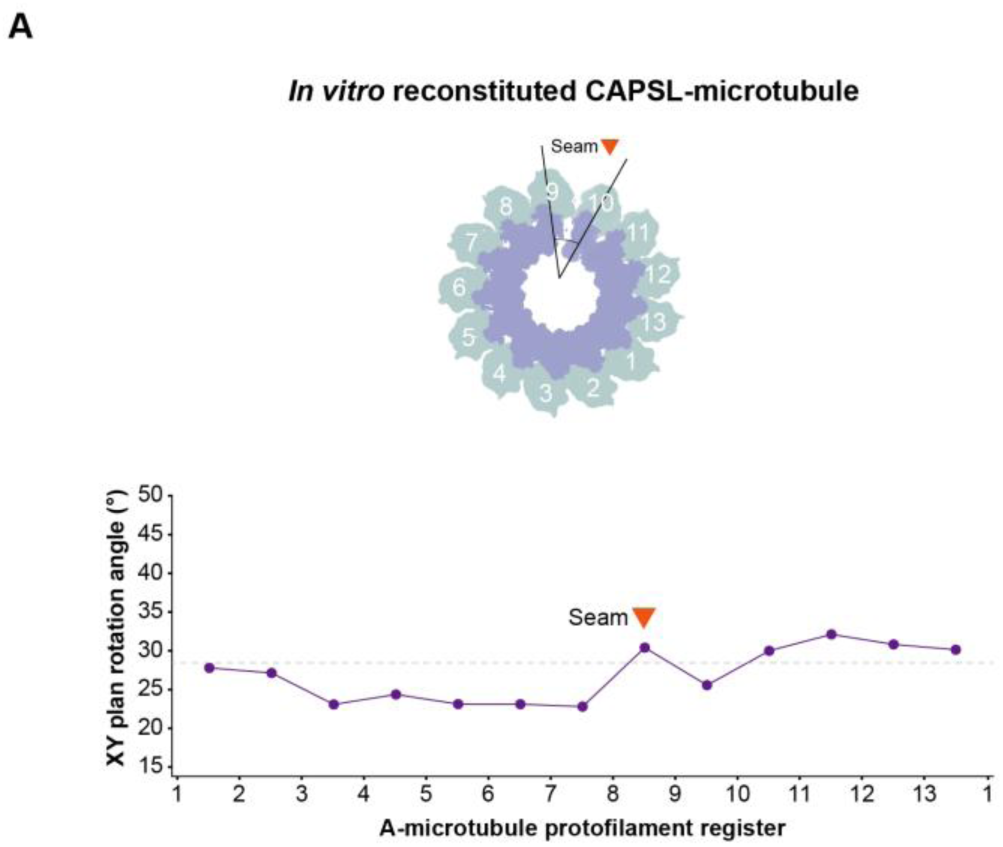
Per-protofilament X-Y plane rotation angles for the *in vitro* CAPSL-microtubule reconstruction, as in Figure 3F-H. The location of the seam is indicated.

**Figure S10.**
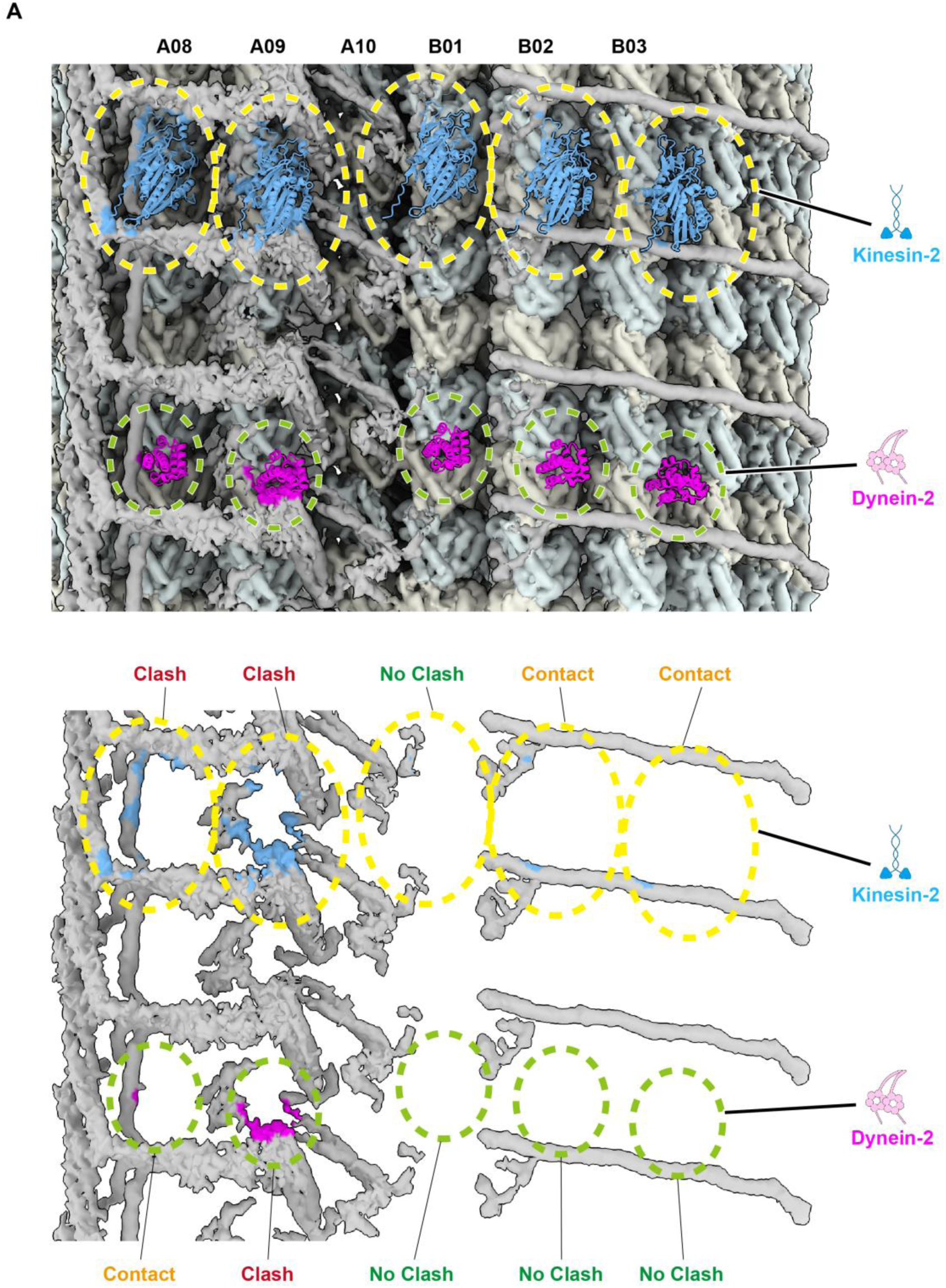
Clashes between kinesin-2 and dynein-2 with transition zone MAPs. **A.** Upper, Docking of multiple AlphaFold predicted complexes formed by tubulin dimer with KIF3A (kinesin-2) and dynein-2 microtubule binding domains into the transition zone doublet reconstruction.Bottom, Clashes between the motor proteins and the transition zone MAPs; blue and purple regions indicate potential clash locations.

**Figure S11.**
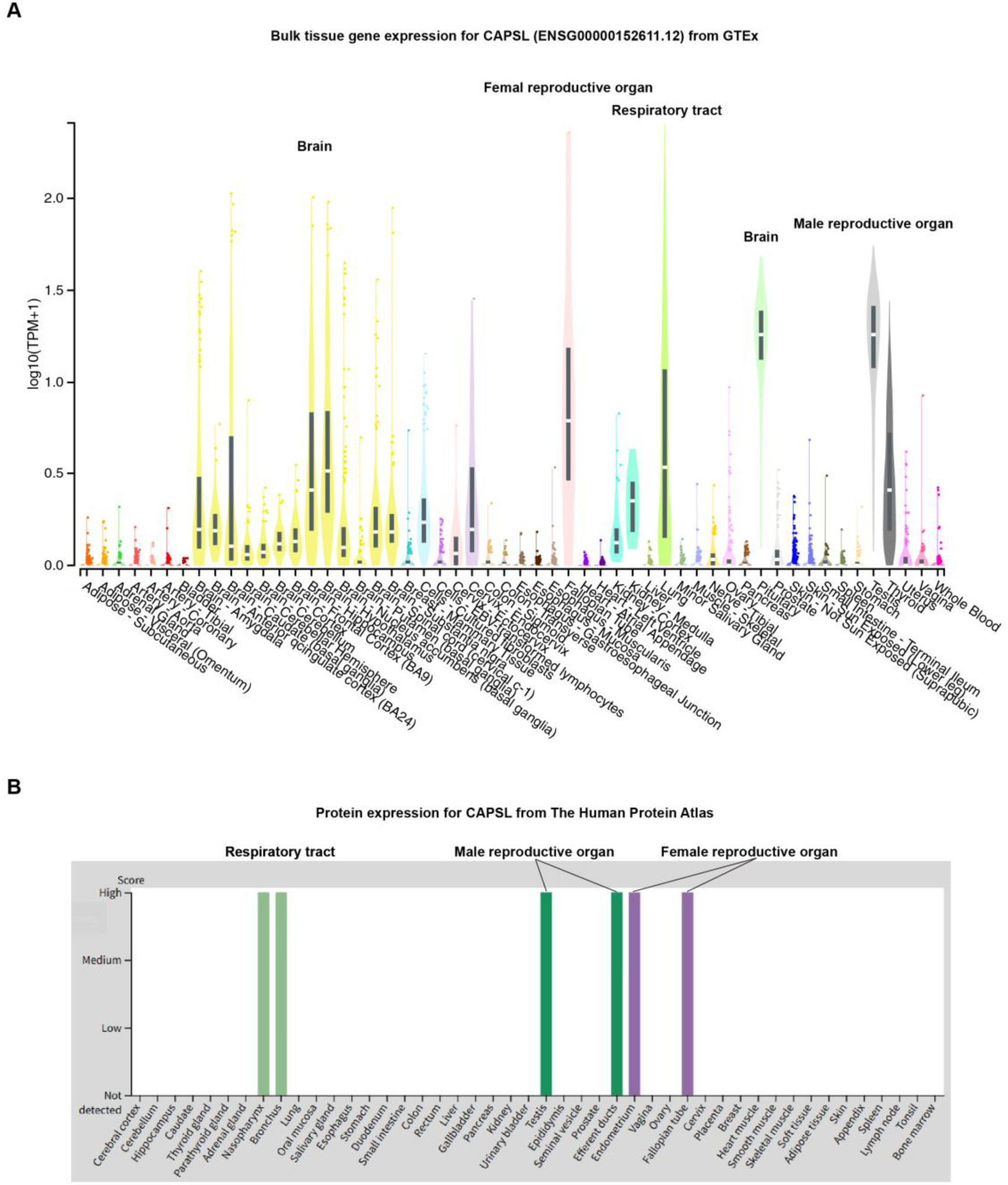
Tissue-dependent expression data for CAPSL. **A.-B.** Tissue-dependent gene expression for CAPSL from the GTEx database (**A**) and the Human Protein Atlas (**B**) (*48, 49*).

**Table S1.** Cryo-ET data collection and model statistics.

|  | Composite map<br>(EMDB-58090)<br>(PDB-3OXP) | A03-A09<br>(EMDB-58085) | Core<br>(EMDB-58086) | B02-B06<br>(EMDB-58087) | B06-B10<br>(EMDB-58088) |
| --- | --- | --- | --- | --- | --- |
| Data collection and processing |  |  |  |  |  |
| Microscope |  | FEI Titan Krios G4 |  | FEI Titan Krios G3i |  |
| Camera |  | Gata K3 BioContinuum GIF |  | Gatan K3 BioQuantum GIF |  |
| Magnification |  | 64,000 |  | 64,000 |  |
| Voltage (kV) |  | 300 |  | 300 |  |
| Electron exposure (e<sup>-</sup>/Å) |  | 160 |  | 160 |  |
| Total tilt series |  | 1,201 |  | 126 |  |
| Defocus range (µm) |  | -2.0 to -4.0 |  | -1.0 to -3.0 |  |
| Pixel size (Å/pixel) |  | 1.384 |  | 1.34 |  |
| Symmetry imposed |  | C1 |  |  |  |
| Initial particle number |  | 680,495 |  |  |  |
| Refinement |  |  |  |  |  |
| Map resolution (Å) | N/A | 4.80 | 4.66 | 4.80 | 4.97 |
| Final particle number | N/A | 51,069 | 51,069 | 51,069 | 51,069 |
| Map sharpening b factor | N/A | -50 | -29 | -53 | -11 |
| Model composition |  |  |  |  |  |
| Atoms | 177,844 |  |  |  |  |
| Protein residues | 36,059 |  |  |  |  |
| Chains | 103 |  |  |  |  |
| R.M.S deviations |  |  |  |  |  |
| Bond length (Å) | 0.002 |  |  |  |  |
| Bond angles (°) | 0.464 |  |  |  |  |
| Validation |  |  |  |  |  |
| MolProbity score | 1.47 |  |  |  |  |
| Clashscore | 3.00 |  |  |  |  |
| Rotamer outlier (%) |  |  |  |  |  |
| Ramachandran plot |  |  |  |  |  |
| Favored (%) | 94.39 |  |  |  |  |
| Allowed (%) | 5.45 |  |  |  |  |
| Outlier (%) | 0.15 |  |  |  |  |
| Masked CC | 0.85 |  |  |  |  |

**Table S2.**
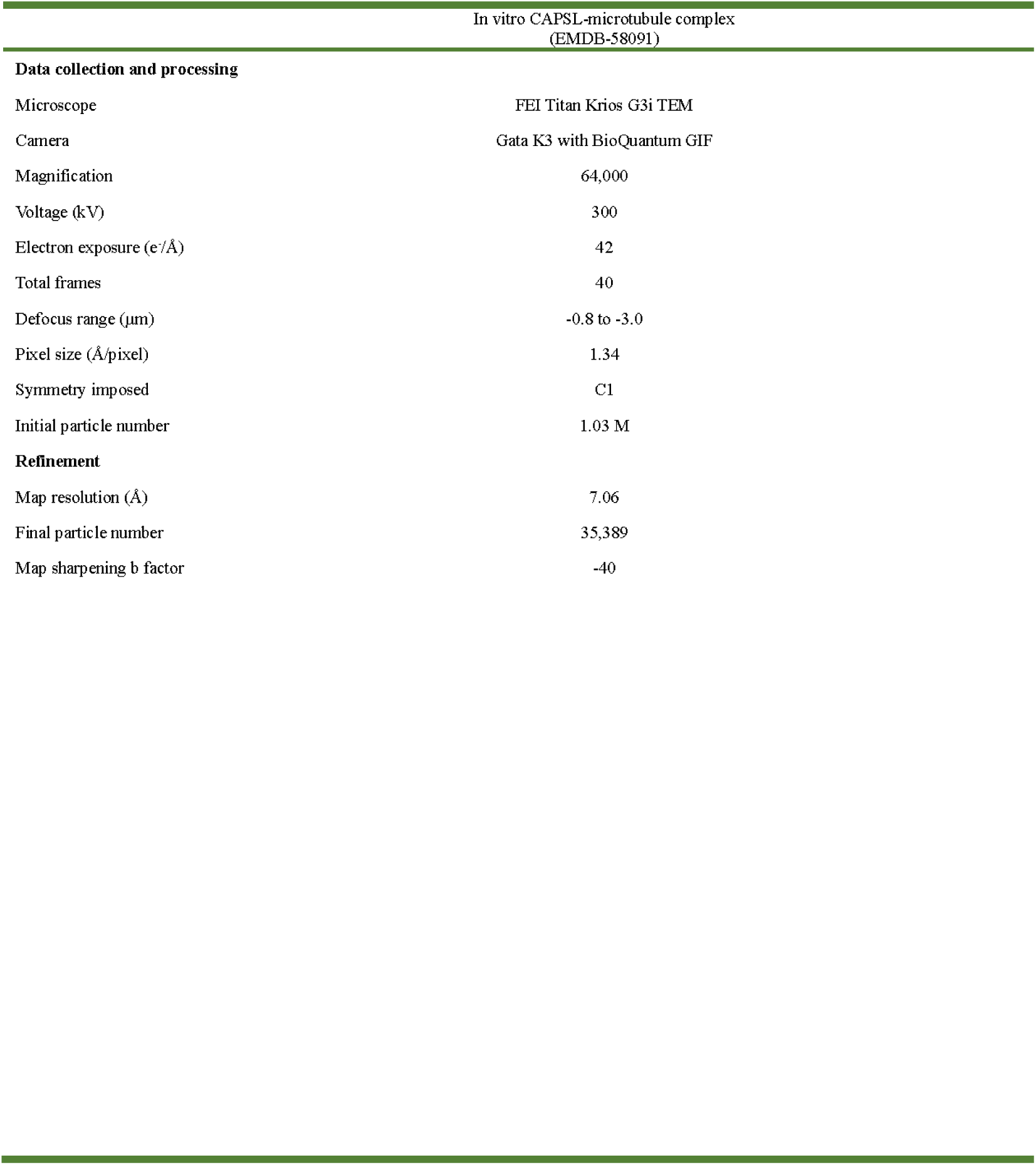
Cryo-EM data collection.

